# Intestinal Serum Amyloid A suppresses systemic neutrophil activation and bactericidal activity in response to microbiota colonization

**DOI:** 10.1101/435503

**Authors:** Caitlin C. Murdoch, Scott T. Espenschied, Molly A. Matty, Olaf Mueller, David M. Tobin, John F. Rawls

## Abstract

The intestinal microbiota influence diverse aspects of host physiology, including the development and function of myeloid lineages. Numerous host and microbial factors are known to poise neutrophils and other granulocytes for response to pathogens and danger signals, yet the mechanisms by which the intestinal microbiota regulate this process are largely unknown. Using gnotobiotic zebrafish, we identified the immune effector Serum amyloid A (Saa) as one of the most highly induced transcripts in digestive tissues following microbiota colonization. Saa is a conserved secreted protein produced in the intestine and liver with described effects on neutrophils *in vitro*, however its *in vivo* functions are poorly defined. We engineered saa mutant zebrafish to test requirements for Saa on innate immunity *in vivo*. Zebrafish mutant for saa displayed impaired neutrophil responses to wounding but augmented clearance of pathogenic bacteria. At baseline, saa mutants exhibited moderate neutrophilia and altered neutrophil tissue distribution. Molecular and functional analyses of isolated neutrophils revealed that Saa suppresses expression of pro-inflammatory mRNAs and bactericidal activity. Saa’s effects on neutrophils depends on microbiota colonization, suggesting this protein mediates the microbiota’s influence on host innate immunity. To test tissue-specific roles of Saa on neutrophil function, we generated transgenic zebrafish over-expressing *saa* in the intestine. Transgenic intestinal saa expression was sufficient to partially complement the neutrophil phenotypes in *saa* mutants. These results indicate Saa produced by the intestine in response to microbiota serves as a systemic signal to neutrophils to restrict aberrant activation, decreasing inflammatory tone and bacterial killing potential while simultaneously enhancing their ability to migrate to wounds.

## Author Summary

The intestine is colonized by dense communities of microorganisms referred to as the microbiota, which impact diverse aspects of host physiology including innate immune system development and function. Neutrophils are phagocytic innate immune cells essential for host defense against infection, and their activity is profoundly impacted by the microbiota although underlying mechanisms remain poorly defined. Here we show the evolutionarily conserved secreted host protein Serum Amyloid A (Saa) mediates microbiota-dependent effects on systemic neutrophil function. Saa is produced by the intestine and liver in response to microbiota, but its *in vivo* functions have remained elusive. Using zebrafish, we demonstrate that Saa promotes neutrophil recruitment to peripheral injury yet restricts clearance of pathogenic bacterial infection. Analysis of isolated neutrophils revealed Saa reduces bactericidal activity and expression of pro-inflammatory genes in a microbiota-dependent manner. Transgenic expression of saa in the intestine of saa mutants was sufficient not only to rescue mutant phenotypes, but also alleviated peripheral wound recruitment defects in germ-free zebrafish. Collectively, these data establish that microbiota-induced Saa in the intestine signals systemically to neutrophils, tuning the extent to which they may be activated by other microbes or respond to injury.

## Introduction

The vertebrate intestine is densely colonized with complex communities of micro-organisms, collectively referred to as the intestinal microbiota. Studies using gnotobiotic animals have demonstrated that microbiota colonization is required for the normal development of an innate immune system capable of mounting appropriate responses to diverse challenges such as infection and injury [1]. Despite being spatially confined to the intestinal lumen by physical and chemical barriers such as the intestinal epithelium and mucus, the microbiota influence both local and systemic host immune development and function [2, 3]. However, the specific mechanisms by which the microbiota impact local and systemic host immune responses remain poorly defined.

Intestinal epithelial cells (IECs) serve as the primary host interface with microbiota and secrete a myriad of factors following microbial colonization [4]. We and others have hypothesized that these microbiota-induced IEC products may mediate the microbiota’s influences on the host immune system [5, 6]. Previous studies have identified a secreted host factor, Serum Amyloid A (Saa), that is potently up-regulated in the intestine following microbial colonization in zebrafish and mice [7–11]. SAA is highly conserved amongst vertebrates, existing as a single gene in fishes and birds and as a multi-gene family in mammals [12, 13]. While basal SAA production is stimulated by the microbiota, SAA production is also markedly augmented following acute injury and infection as a part of the acute phase response, whereby circulating levels can reach 1 mg/mL [13–15]. Moreover, SAA is elevated in chronic pathological conditions where both local and circulating concentrations are positively correlated with inflammation. Accordingly, SAA is an established biomarker for chronic inflammatory diseases such as diabetes, atherosclerosis, and inflammatory bowel disease (IBD) [16–20]. Taken together, SAA’s high degree of evolutionary conservation coupled with its strong induction following inflammatory stimuli suggests important roles for SAA in health and disease.

Previous studies have reported both pro-and anti-inflammatory effects of SAA on host immune responses. The vast majority of these studies have been conducted *in vitro* using recombinant human SAA (rhSAA), which has been shown to directly influence granulocytes, including monocytes and neutrophils, promoting the production of inflammatory cytokines, reactive oxygen species (ROS), and directing motility [21–25]. Recent reports have also shown that mammalian SAAs can bind retinol and mediate host responses during infection [8]. Moreover, induction of SAA in the intestine following colonization with specific microorganisms such as segmented filamentous bacteria (SFB) can shape local adaptive immune cell development by promoting Th17 differentiation [26, 27]. However, a fuller assessment of SAA’s *in vivo* functional roles has remained elusive due to the existence of multiple SAA gene paralogs in mammals (3 in humans, 4 in mice) and the use of cell-culture based assays performed with rhSAA that behaves dissimilarly to endogenous SAA protein [28, 29].

The existence of a single SAA ortholog in fishes provides interesting opportunities to define SAA’s *in vivo* functional roles. We previously demonstrated that partial (~30%) knockdown of *saa* transcript in zebrafish influenced tissue-specific neutrophil behaviors *in vivo*, leading us to hypothesize that Saa regulates neutrophil activity in homeostasis [12]. However, Saa’s influence on systemic neutrophil activation and function in homeostasis and in biologically relevant contexts such as injury and infection remained unresolved. Neutrophils are professional phagocytic myeloid cells that play critical roles in host defense against pathogens. The most abundant immune cell in circulation and the first to be recruited to sites of injury, neutrophils eliminate microbial invaders and debris through a variety of mechanisms including phagocytosis, generation of ROS, and secretion of anti-microbial peptides [30, 31]. Neutrophils are conditioned by host and microbially derived signals, including pathogen associated molecular patterns (PAMPs) and damage associated molecular patterns (DAMPs), allowing for proper responses to inflammatory stimuli [32, 33]. Previous studies have shown that microbial colonization of the intestine promotes neutrophil differentiation, activation, and response to peripheral injury [10, 12, 34–39]. However the signaling molecules that mediate these interactions *in vivo* remain largely unknown.

Here, using zebrafish, we demonstrate that Saa is a host factor that signals microbial status in the intestine to extra-intestinal populations of immune cells and directs their responses to inflammatory stimuli. Zebrafish share highly conserved hematopoietic programs with other vertebrates, including specification of myeloid lineages, which can be coupled with optical transparency and genetic tractability to allow for high resolution *in vivo* imaging of innate immune processes [40, 41]. Moreover, the zebrafish genome encodes a single *saa* ortholog, allowing us to generate the first-ever *saa* null vertebrate model. By comparing wild-type zebrafish to those that lack *saa* or express it only in intestinal epithelial cells under conventional and gnotobiotic conditions, we reveal Saa’s impact on systemic neutrophil activity in homeostasis and following bacterial infection and wounding. Using tissue specific rescue, we demonstrate that intestinally-derived Saa can shape systemic neutrophil function and can even restore neutrophil defects observed in germ-free zebrafish.

## Results

### Saa promotes neutrophil migration during injury and homeostasis

To investigate Saa’s effects on neutrophil function *in vivo*, we first generated *saa* mutant zebrafish, identifying three independent deletion alleles all resulting in frameshift mutations within *saa* exon 2. The largest *saa* deletion allele (22 bp, designated *rdu60,* homozygous mutants hereafter referred to as *saa*^−/−^) resulted in 90% reduced *saa* mRNA (S1 Fig A-E). *saa*^−/−^ zebrafish survive to adulthood and exhibit no gross developmental abnormalities (S1 Fig F-J). We performed caudal fin amputations on WT and *saa*^−/−^ *Tg(lyz:EGFP)* larvae and quantified neutrophil recruitment to the wound margin, observing fewer neutrophils at 6 hours post-wounding in *saa*^−/−^ larvae (Fig 1A,B). Further, *in vivo* imaging revealed that neutrophils in the vicinity of the wound moved with slower mean velocity in *saa* mutant larvae (Fig 1C). Importantly, *saa* mRNA was not upregulated at 6 hours post amputation in WT larvae (S1 Fig L), demonstrating acute *saa* induction does not affect neutrophil activity. Thus, Saa is required for neutrophil mobilization to sites of injury independent of systemic induction.

**Fig 1.**
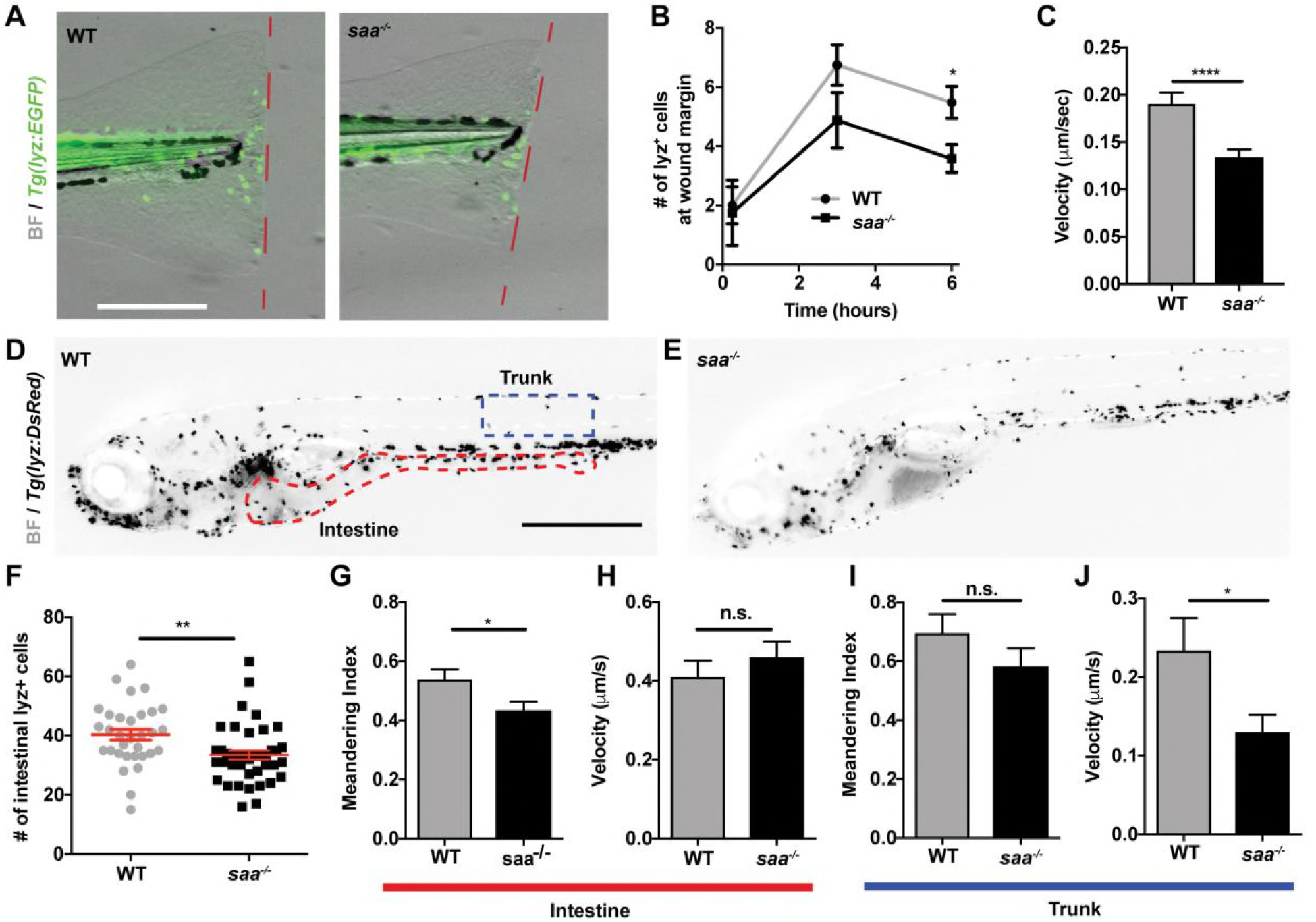
Saa mediates neutrophil behavior *in vivo*. **(A-B)** Imaging and quantification of lyz:EGFP+ neutrophils recruited to tail wound margin over 6 hours following caudal fin amputation (dashed red line indicates wound margin; scale bar = 250 μm) (n ≥ 24 larvae / genotype at 6 hour time point). **(C)** Measurement of lyz:DsRed+ neutrophil velocity from time-lapse imaging in caudal fin tissue over 6 hour period following amputation (n = 4 larvae / genotype, 87-112 cells tracked / genotype). **(D-E)** Representative images of 6 dpf *Tg(lyz:DsRed)* WT and *saa*−/− larvae (scale bar = 500 μm). **(F)** Enumeration of intestine-associated lyz:EGFP+ cells in 6 dpf larvae (n = 32-40 larvae / genotype). **(G-J)** Quantitative analysis of lyz:EGFP+ neutrophil behavior from time-lapse imaging of distinct anatomical compartments (intestine and trunk, ROIs in panel D) in 6 dpf larval zebrafish (6 larvae / genotype, ≥ 23 cells analyzed / genotype / tissue). Data analyzed by *t*-test. For panel B, statistical comparisons were performed within each time point. Data are presented as mean ± SEM. * *p* < 0.05, ** *p* < 0.01, *** *p* < 0.001, **** *p* < 0.0001

Given that Saa affects neutrophil responses to injury, we tested if Saa regulates basal neutrophil behavior. Analysis of neutrophils in homeostasis revealed Saa promotes neutrophil velocity in the trunk and linear migration in the intestine (Fig 1G-J), consistent with our prior *saa* morpholino data [12]. Considering *saa* is expressed in IECs [10], we reasoned that Saa may promote neutrophil recruitment to the intestine. Indeed, we observed fewer intestine-associated neutrophils in *saa*^−/−^ larvae (Fig 1D-F). These data demonstrate that Saa promotes neutrophil recruitment to peripheral injury and to distinct tissues during development.

### Saa restricts systemic neutrophil abundance and bactericidal activity

Given that *saa* loss is associated with impaired neutrophil recruitment to wounds and healthy tissues, we asked whether systemic neutrophil abundance is altered in *saa* mutants. We enumerated systemic neutrophils by flow cytometry (Fig 2A, S2 Fig A) and observed elevated neutrophil abundance in *saa*^−/−^ larvae. This was corroborated by increased expression of the granulocyte marker genes *lysozyme C (lyz)*, *l-plastin (lcp)* and the granulopoetic cytokine *colony stimulating factor 3a* (*csf3a*, also known as *gcsf1a*) in 6 days post fertilization (dpf) whole larvae (Fig 2D-F). Morphological classification of lyz^+^ neutrophils into sub-populations from cytospin preparations (adapted from [42] revealed an over-representation of immature lyz^+^ neutrophils in *saa* deficient animals compared to WT controls (Fig 2B,C). Together, these data demonstrate a novel role for Saa regulating neutrophil maturation *in vivo*.

**Fig 2.**
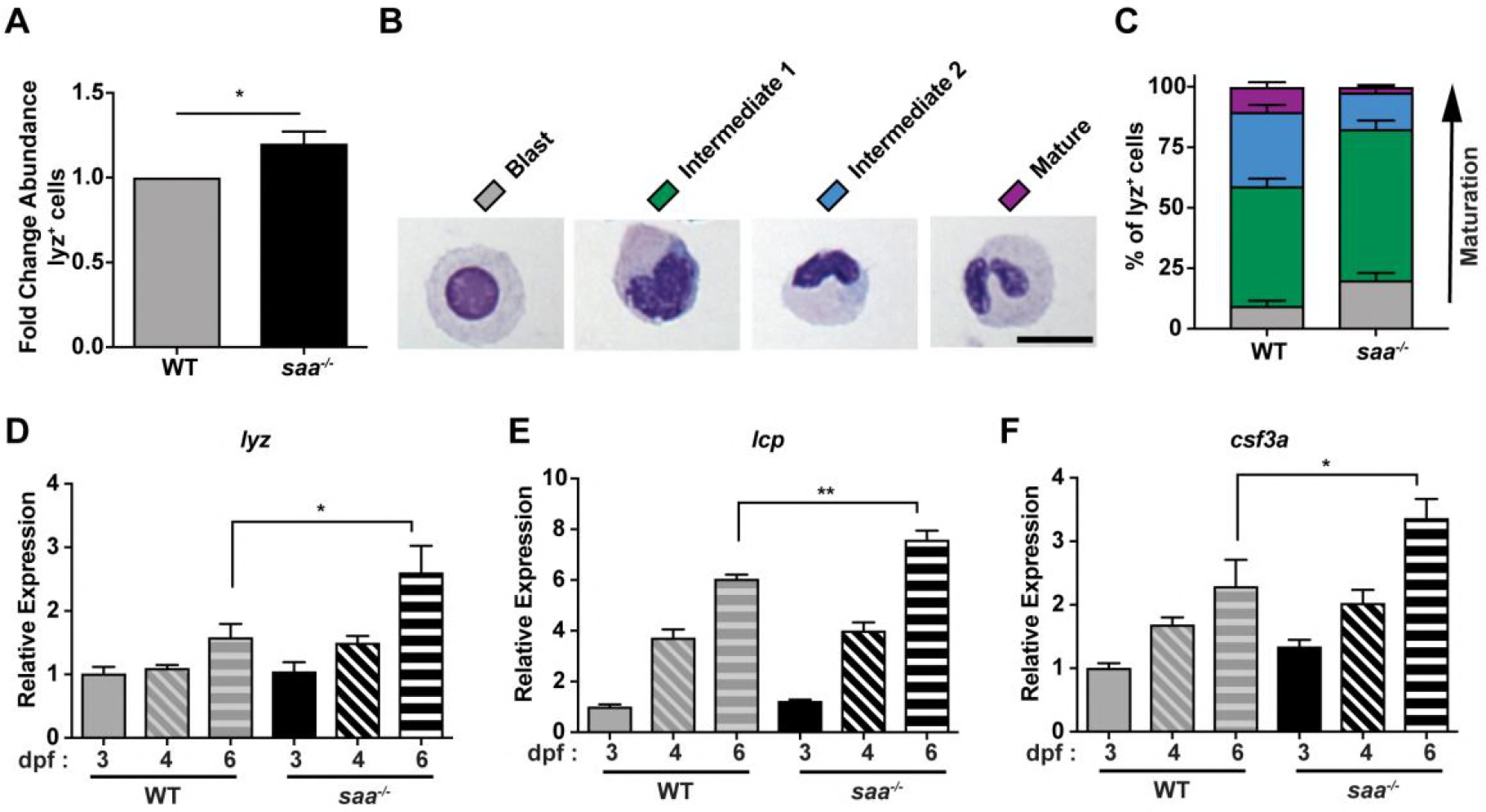
Saa regulates neutrophil abundance and maturation. **(A)** Flow cytometry analysis of lyz:EGFP+ neutrophil abundance from whole 6 dpf WT and *saa* mutant zebrafish larvae (results are combined from 3 independent experiments, ≥ 4 replicates / genotype / experiment, 60-90 larvae / replicate). **(B-C)** Morphological analysis of lyz:DsRed+ neutrophil cytospins stained with Wright-Giemsa from adult WT and *saa* mutant zebrafish kidneys (5-6 adult zebrafish kidneys / genotype / experiment, 2 independent experiments, n ≥ 199 cells analyzed / genotype / experiment) (scale bar = 10 μm). **(D-F)** qRT-PCR of leukocyte-associated transcripts *l-plastin (lcp), lysozyme C (lyz),* and *colony-stimulating factor 3a (csf3a)* from 6 dpf whole zebrafish larvae (n = 4 replicates / genotype / timepoint, 25-30 larvae / replicate). Data in panel A analyzed by *t*-test. Data in panel C analyzed by chi-squared test. Data in panels D-F analyzed by one-way ANOVA with Tukey’s multiple comparisons test. Data are presented as mean ± SEM. * *p* < 0.05, ** *p* < 0.01, *** *p* < 0.001, **** *p* < 0.0001

Exposure to MAMPs or inflammatory host molecules can elicit defined neutrophil transcriptional responses, reflecting their activation state [43–47]. Gene expression analysis of FACS-isolated neutrophils revealed elevated expression of genes encoding pro-inflammatory cytokines (*tnfa*, *il1b*), antimicrobial peptides (*pglyrp2*, *pglyrp5*), and regulators of ROS production (*mpx*, *ncf1*) in *saa*^−/−^ larvae (Fig 3A). These transcriptional differences suggest Saa restricts basal neutrophil activation. As neutrophils primarily function to clear microbial infections [48, 49], we co-cultured adult zebrafish kidney neutrophils with *Escherichia coli* then assessed bacterial viability. Isolated neutrophils from both WT and *saa*^−/−^ fish were viable *ex vivo* and exhibited morphological responses (e.g., extending cytosolic projections) to bacteria (Fig 3B, S3 Fig A-E,J). Co-culture with *E. coli* induced *il1b* mRNA in WT neutrophils, demonstrating zebrafish neutrophils respond transcriptionally to bacteria *ex vivo* (Fig 3C). Moreover, *il1b* transcript levels were significantly increased in unstimulated *saa* mutant adult kidney neutrophils compared to WT, consistent with our observations from larval neutrophils (Fig 3A,C). However, following co-culture with *E. coli*, *il1b* in WT neutrophils reached similar levels measured in *saa*^−/−^ neutrophils (Fig 3C).

**Fig 3.**
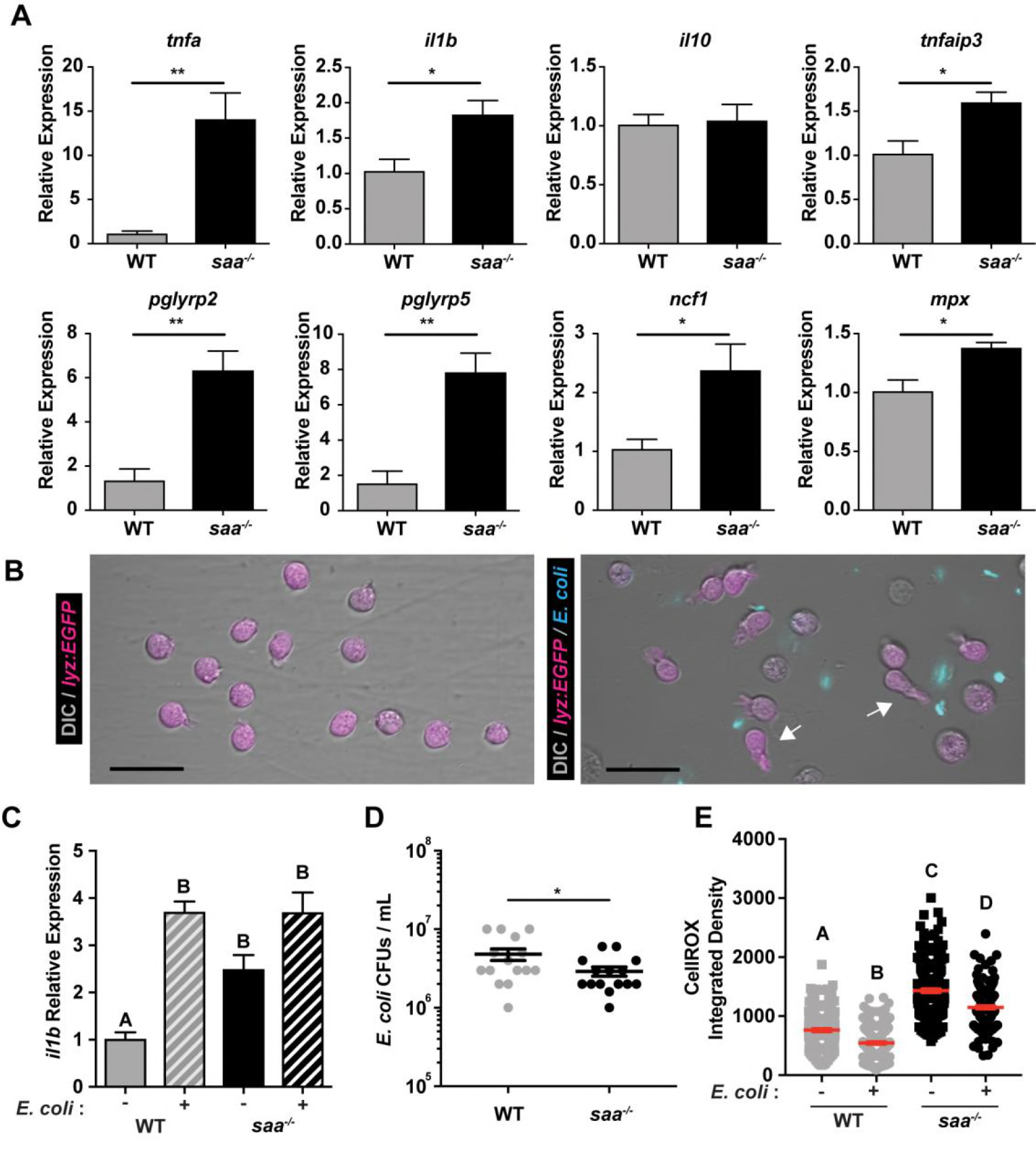
Saa suppresses neutrophil transcriptional activation and bactericidal activity. **(A)** qRT-PCR of pro-inflammatory mRNAs from sorted neutrophils from 6 dpf WT and *saa* mutant zebrafish larvae (5,000-12,000 lyz:EGFP+ cells / replicate, 3-6 replicates / genotype / experiment, 60-90 larvae / replicate). **(B)** Microscopic analysis shows neutrophils isolated from adult zebrafish kidneys extend protrusions in response to bacterial signals *ex vivo* (white arrows; scale bar = 20 μm). **(C)** *il1b* expression in un-stimulated and *E. coli* exposed lyz:EGFP+ neutrophils from WT and *saa* mutant zebrafish following 4 hours *ex vivo* culture (3-5 replicates / genotype / condition). **(D)** CFU quantification of bacterial concentration following 4 hour co-culture of isolated lyz:EGFP+ neutrophils from WT and *saa* mutant zebrafish with *E. coli* (MOI 2). **(E)** Quantification of intracellular ROS levels by CellROX fluorescence from neutrophils cultured *ex vivo* with and without *E. coli* (lyz:EGFP+ cells isolated from 6 zebrafish adult kidneys / genotype, quantification of ≥ 177 cells / condition). In panels C and E, a one-way ANOVA with Tukey’s multiple comparisons test was used. Data in panel D was analyzed with a *t-*test. Data are presented as mean ± SEM. * *p* < 0.05, ** *p* < 0.01, *** *p* < 0.001, **** *p* < 0.0001

To assess neutrophil bactericidal activity, we enumerated CFUs following 4 hours of co-culture and found *saa*^−/−^ neutrophils killed significantly more bacteria than WT neutrophils (Fig 3D). To interrogate possible mechanisms of bacterial clearance, we labeled neutrophils with CellROX *ex vivo* and measured levels of intracellular ROS by confocal microscopy. Bacterial exposure resulted in decreased ROS in both WT and *saa*^−/−^ neutrophils, indicating bacteria stimulate neutrophil degranulation. Interestingly, neutrophils from *saa* mutant animals had elevated levels of ROS relative to WT controls both at baseline and after bacterial stimulation (Fig 3E), which is consistent with their augmented bacterial killing activity (Fig 3D). Collectively, these data indicate neutrophils from *saa* mutants are aberrantly activated as evidenced by elevated pro-inflammatory mRNA expression, augmented bactericidal activity, and elevated ROS production, and suggest Saa restricts systemic neutrophil inflammatory tone *in vivo*.

### Intestinally-derived Saa alters neutrophil distribution and activity

Since *saa* is highly expressed in the larval zebrafish intestine and liver [10], it is possible that Saa produced by specific tissues differentially affects systemic neutrophil conditioning. Given that the intestine serves as a primary interface between the host and microbiota, and that *saa* is transcriptionally upregulated in the intestine following microbial colonization, we engineered transgenic zebrafish in which *saa* is expressed specifically in IECs. While several zebrafish promoters have been identified with intestine-restricted activity, they only drive transgene expression in subsets of IECs (e.g., the *fabp2*/*ifabp* promoter is active in anterior enterocytes) [10, 50]. Since *cldn15la* is expressed broadly in IECs [51, 52] we queried adult zebrafish IEC FAIRE-seq data and identified a 349 bp open chromatin region in the *cldn15la* promoter [53]. This element contains predicted binding sites for intestine-specific transcription factors such as Cdx2, and appears to drive expression in all IECs (S4 Fig A-H) [53, 54]. We generated *Tg*(*-0.349cldn15la:saa;cmlc2:EGFP*) zebrafish [subsequently denoted *Tg(cldn15la:saa)*], which express full-length zebrafish *saa* in IECs (S4 Fig I-J).

We crossed this transgene into the *saa* mutant background, and asked whether IEC-derived Saa could complement *saa*^−/−^ neutrophil defects. As expected, we observed fewer intestine-associated neutrophils in each intestinal segment of *saa*^−/−^ larvae (from anterior to posterior, segments 1-3). Intestinally-derived Saa was sufficient to complement this mutant phenotype, restoring intestine-associated neutrophil numbers to WT levels (Fig 4A,B). Thus, Saa produced in IECs is sufficient to promote neutrophil recruitment to the intestine, suggesting Saa is a neutrophil chemoattractant *in vivo*. To determine if neutrophil function was altered by intestinally-derived Saa, we performed caudal fin amputations. At 6 hours post amputation, neutrophil recruitment to the wound in *saa*^−/−^*;Tg(cldn15la:saa)* larvae was equivalent to the WT response (Fig 4C), demonstrating intestinally-derived *saa* is sufficient to restore neutrophil mobilization to the caudal fin in in otherwise *saa* deficient larvae.

**Fig 4.**
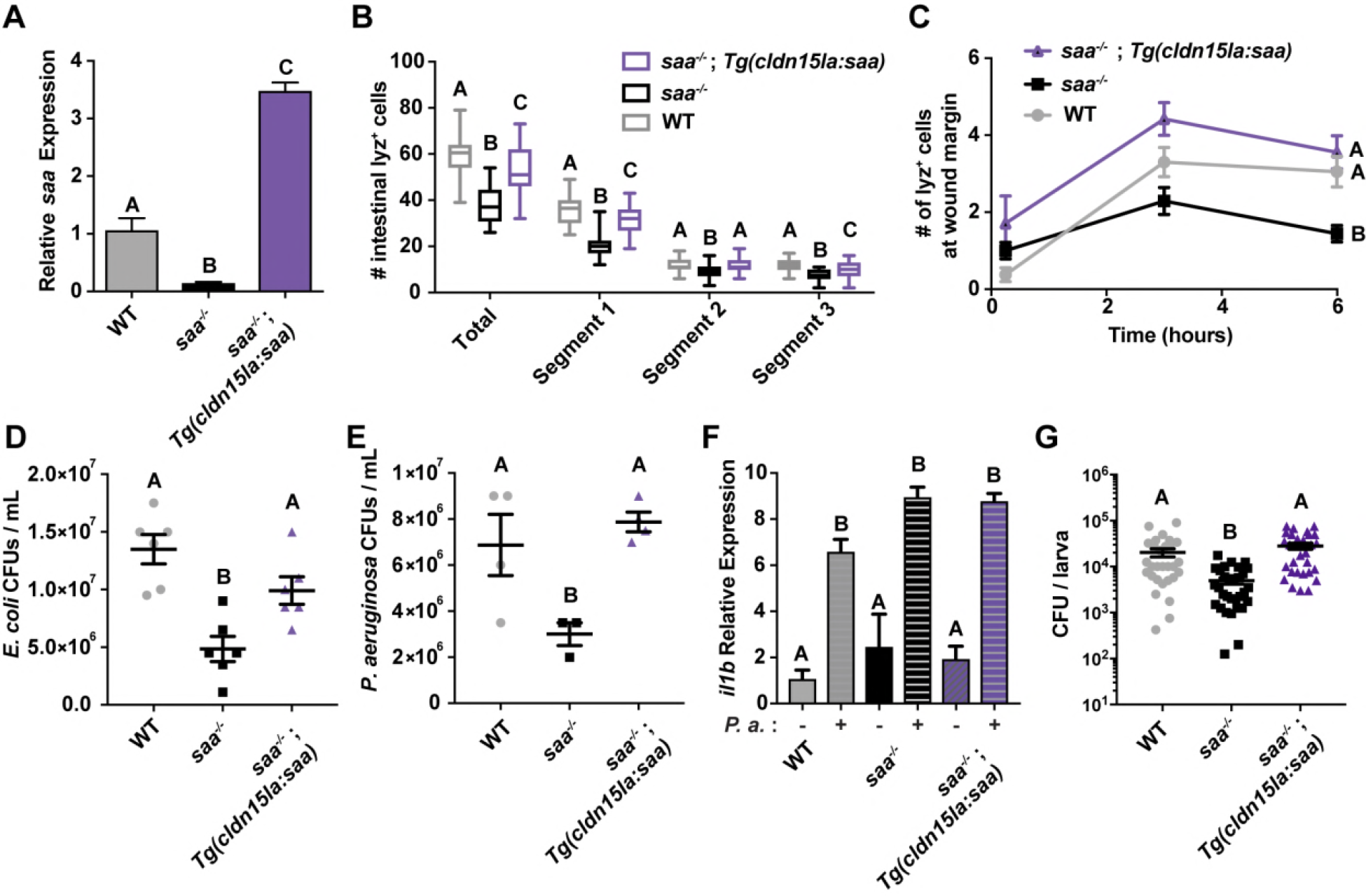
Intestinally-derived Saa regulates systemic neutrophil activity. **(A)** qRT-PCR of *saa* from whole 6 dpf larvae of the indicated genotypes (n = 4 replicates / genotype, 25-30 larvae / replicate) **(B)** Enumeration of intestine-associated lyz:DsRed^+^ neutrophils along the anterior to posterior axis (segment 1 to segment 3) in 6 dpf larvae (n = 30 larvae / genotype). **(C)** lyz:DsRed^+^ neutrophil recruitment to caudal fin wound 6 hours following amputation in 6 dpf zebrafish larvae (n ≥ 18 larvae / genotype at 6 hour time point). **(D-E)** CFU quantification of bacterial concentration following 4 hour co-culture of lyz:DsRed^+^ adult zebrafish neutrophils with both *E. coli* (D, MOI 2) and *P. aeruginosa* (E, *P.a.*, MOI 0.2) (3-6 replicates / genotype). **(F)** *il1b* qRT-PCR from lyz:DsRed^+^ neutrophils co-cultured with and without *P.a. ex vivo* for 4 hours (n ≥ 2 replicates / condition). **(G)** CFU quantification of *in vivo P.a.* bacterial burden following systemic infection of larval zebrafish at 5 days post infection (dpi) (data from 3 independent experiments, n ≥ 30 larvae / genotype). Data in panels A-G were analyzed by one-way ANOVA with Tukey’s multiple comparisons test. For panel C, statistical comparisons were performed amongst samples within the same time point. Data are presented as mean ± SEM. * *p* < 0.05, ** *p* < 0.01, *** *p* < 0.001, **** *p* < 0.0001

To further investigate the effects of intestinally-derived Saa on neutrophil function, we quantified neutrophil bactericidal activity from WT, *saa*^−/−^, and *saa*^−/−^*;Tg(cldn15la:saa)* zebrafish. We used *E. coli* which is cleared quickly by the zebrafish immune system and *Pseudomonas aeruginosa* which is capable of establishing systemic infections [55, 56]. We observed reduced survival of both *E. coli* and *P. aeruginosa* following co-culture with *saa*^−/−^ neutrophils vs WT controls. However, neutrophils from *Tg(cldn15la:saa)*^+^ *saa*^−/−^ zebrafish exhibited comparable bactericidal activity to WT neutrophils (Fig 4D,E). These findings demonstrate intestinally-derived Saa is sufficient to constrain bactericidal activity of kidney neutrophils. Considering the profound effects of *saa* levels and source on neutrophil antibacterial function *ex vivo*, we asked if Saa influenced bacterial clearance *in vivo* using systemic *P. aeruginosa* infection. Consistent with our *ex vivo* results, we observed enhanced bacterial clearance in *saa*^−/−^ larvae which is returned to WT levels in *saa*^−/−^*;Tg(cldn15la:saa)* larvae (Fig 4G).

Enumeration of systemic neutrophil abundance by flow cytometry in *saa*^−/−^*;Tg(cldn15la:saa)* revealed that intestinally-derived Saa failed to alleviate the neutrophilia observed in *saa*^−/−^ larvae (S5 Fig A). Moreover, qRT-PCR analysis of neutrophils isolated from *saa*^−/−^*;Tg(cldn15la:saa)* larvae revealed persistently elevated expression of pro-inflammatory and anti-microbial effectors (*pglyrp2*, *tnfa*, *il1b*, *ncf1*) (S5 Fig B). Thus, intestinal expression of *saa* was sufficient to rescue only a subset of neutrophil defects observed in *saa* mutant animals. These observations highlight the potential requirement for other tissue sources or temporal control of Saa to condition different aspects of systemic neutrophil function. Collectively, these data demonstrate that expression of *saa* in the intestinal epithelium is sufficient to promote neutrophil recruitment to a peripheral wound and to restrict bactericidal activity *in vivo* and *ex vivo*, but is unable to dampen neutrophilia or elevated neutrophil pro-inflammatory mRNA profiles observed in *saa* mutant zebrafish.

### Microbiota-induced Saa suppresses neutrophil pro-inflammatory mRNA production and promotes migration to a tail wound

We and others have shown that intestinal microbiota influence a variety of neutrophil phenotypes both in homeostasis and following injury [12, 36–38]. Since our results indicate that Saa suppresses neutrophil activation (Fig 3A,E) and previous studies have reported Saa has direct bactericidal activity [57–59] we speculated that *saa* deficiency may impact microbiota composition. We used 16S rRNA gene sequencing to compare intestinal bacterial communities in the digestive tracts of *saa*^−/−^ and WT sibling zebrafish at both larval and adult stages. We found that *saa* genotype had no significant effects on gut bacterial community composition as measured by alpha-or beta-diversity metrics at larval (6 dpf) or adult (70 dpf) stages (S2 and S3 Tables). These findings demonstrate that Saa does not broadly impact gut microbiota composition in zebrafish.

Since *saa* is potently induced following microbial colonization (Fig 5A), we asked if Saa regulates neutrophil activation in response to the microbiota. We isolated neutrophils from 6 dpf gnotobiotic WT and *saa*^−/−^ larvae, and found expression of pro-inflammatory mRNAs was significantly elevated in WT neutrophils from conventionalized (CV) zebrafish vs germ-free (GF) controls, confirming microbiota-derived signals induce neutrophil pro-inflammatory mRNA expression (Fig 5B). Comparison of the same transcripts in neutrophils from *saa*^−/−^ zebrafish reared under CV and GF conditions revealed augmented induction of mRNAs following microbiota colonization. Since pro-inflammatory mRNA expressions was comparable between neutrophils from WT GF and *saa*^−/−^ GF larvae, we conclude the microbiota potentiate transcriptional activation observed in *saa*^−/−^ larvae (Fig 5B). These data demonstrate that Saa functions to restrict aberrant activation of neutrophils by the microbiota at homeostasis.

**Fig 5.**
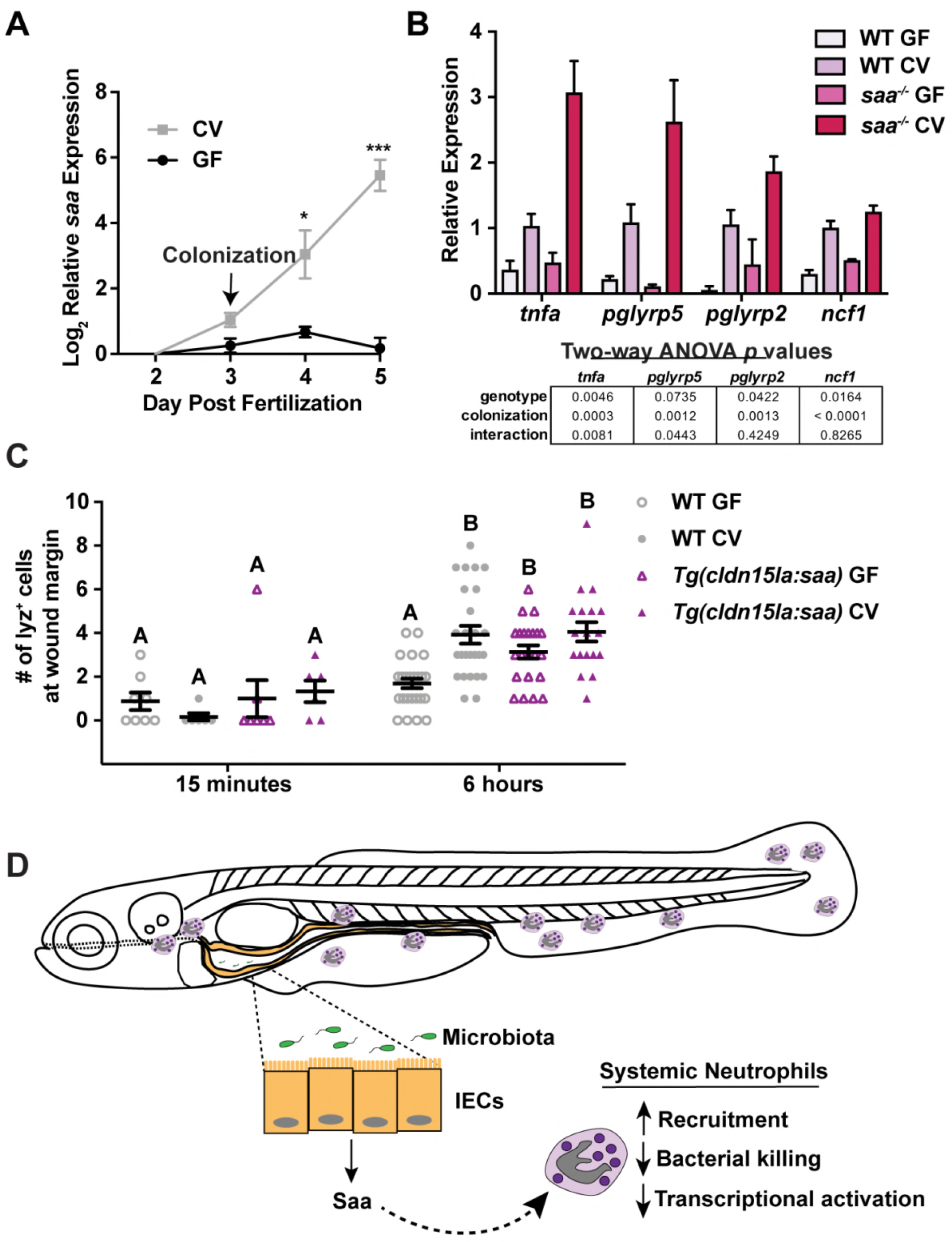
Microbiota-induced Saa conditions neutrophils *in vivo*. **(A)** qRT-PCR of *saa* from gnotobiotic zebrafish larvae following microbiota colonization (CV) at 3 dpf versus germ free (GF) (3 replicates / condition / timepoint, n ≥ 20 larvae / replicate). **(B)** qRT-PCR of neutrophils isolated from 6 dpf gnotobiotic WT and *saa* mutant zebrafish larvae (3 replicates / genotype / condition, n ≥ 27 larvae / replicate). **(C)** lyz:EGFP^+^ neutrophil recruitment to caudal fin wound margin 6 hours after amputation in 6 dpf gnotobiotic WT and *Tg(cldn15la:saa)* sibling zebrafish larvae (n ≥ 18 larvae / genotype / condition at the 6 hour timepoint). **(D)** Proposed model of systemic neutrophil conditioning by intestinal Saa. Statistical comparisons of data in panel in A were performed within each time point and analyzed by *t*-test. Data in panels B and D analyzed by one-way ANOVA with Tukey’s multiple comparisons test. Data in panel C analyzed by two-way ANOVA with *p* values reported in the table. Data are presented as mean ± SEM. * *p* < 0.05, ** *p* < 0.01, *** *p* < 0.001, **** *p* < 0.0001

Given that the microbiota induce *saa* in the intestine, and germ-free zebrafish exhibit neutrophil defects [10], we asked if transgenic *saa* expression in IECs was sufficient to rescue neutrophil deficiencies in GF animals. Consistent with previous findings, we observed reduced neutrophil wound recruitment in GF WT larvae vs CV controls [12]. Transgenic intestinal *saa* expression in GF larvae rescued neutrophil mobilization to the wound margin 6 hours after injury (Fig 5C). These data support a model wherein Saa produced by IECs in response to the microbiota limits aberrant neutrophil activation by the microbiota, allowing neutrophils to effectively respond to challenges such as peripheral injury (Fig 5D).

## Discussion

Previous reports in mice and zebrafish demonstrated that microbiota influences diverse aspects of neutrophil biology, including increased abundance and longevity, enhanced wound recruitment, and elevated bacterial killing[10, 12, 34–36, 38, 39, 60]. However, few studies conclusively identify host or microbial factors that mediate the microbiota’s influence on neutrophils. We demonstrated that Saa induced following colonization transduces information regarding the intestinal microbiota, regulating diverse aspects of neutrophil biology. In colonized zebrafish at homeostasis, Saa promoted neutrophil maturation and mobilization to the intestine while suppressing systemic neutrophil abundance and pro-inflammatory gene expression. During inflammatory challenge, Saa restricted neutrophil anti-bacterial activity yet promoted neutrophil recruitment to a wound. While *in vitro* studies suggest rhSAA promotes pro-inflammatory cytokine and ROS production (TNFα, IL-1β, and IL-8)[23, 61–65], the pro-inflammatory effects of rhSAA (rhSAA; Peprotech, Rocky Hill, NJ, USA) on granulocytes contrasts with reports using purified endogenous SAA [28, 29]. Thus, our *in vivo* analysis clarifies conflicting reports of Saa’s effects on neutrophils.

Our results establish Saa as a major host factor mediating microbial control of neutrophil function. As *saa* is potently induced following microbial colonization, we reasoned that neutrophil defects in *saa*^−/−^ larvae would phenocopy those exhibited in WT germ-free larvae. Supporting this hypothesis, we observed decreases in neutrophil migratory behavior, intestinal association, and wound recruitment in GF WT and colonized *saa*^−/−^ larvae [10, 12, 66, 67]. We demonstrated intestinally-derived Saa was sufficient to partially complement neutrophil deficiencies in *saa*^−/−^ zebrafish and could restore neutrophil wound recruitment in defects GF larvae. These results indicate that intestinally-derived Saa conditions neutrophils *in vivo* following microbiota colonization. As we have previously shown that colonization with different microbial taxa leads to varied levels of *saa* induction [11], this signaling axis may facilitate taxa-specific effects on host innate immune development. Further, it is possible that mammalian SAA paralogs similarly condition neutrophils, as a subset of *Saa* genes are induced in mouse intestinal tissues following microbial colonization [7–9, 57]. As microbiota-stimulated intestinal Saa production regulates systemic immune cell function, broad-spectrum antibiotics usage can dramatically impact the microbiota [68–70], and antibiotic treatment results in reduced mouse intestinal *Saa* [71], our model predicts antibiotic treatment would be associated with aberrant neutrophil function. We speculate that secondary infections that can occur following antibiotic use [72] could be due in part to concomitant alterations in SAA production.

Our discovery that Saa functions in the microbiota-neutrophil axis motivates interest in underlying mechanisms. Neutrophil functions are largely regulated through “priming”, whereby microbial products (e.g. LPS, peptidoglycan, and flagellin) and host factors (e.g. TNFα, IL1-β, IL-8, GM-CSF, and ATP) signal to neutrophils, preparing them for response to additional stimuli [32, 73]. Intriguingly, many priming factors are induced in the zebrafish and mammalian intestine by the microbiota (e.g. TNFα, IL1-β, and GM-CSF) [7, 8, 10, 53, 74]. Primed neutrophils exhibit enhanced bacterial killing, altered motility, and transcriptional changes [33]. Once believed to be transcriptionally quiescent, several studies have reported neutrophil transcriptional responses to various priming stimuli *in vitro* [43–45, 47] and *in vivo* [46, 75, 76]. We demonstrate Saa restricts neutrophil pro-inflammatory gene expression and show induction of pro-inflammatory genes in neutrophils from colonized vs GF WT zebrafish, which is augmented in *saa*^−/−^ larvae. Consistent with our gene expression results, Saa limits primed neutrophil phenotypes of ROS production and bacterial killing *in vivo* and *ex vivo*. We propose Saa is upregulated following microbiota colonization to temper aberrant neutrophil priming by microbial products and production of host inflammatory mediators, thus limiting collateral damage to host tissues [77–79].

Our data reveal that Saa promotes neutrophil maturation in adult zebrafish. A recent study showed that mature neutrophils exhibit increased motility and response to stimuli [80]. While it is possible that altered neutrophil maturation may underlie the phenotypes we observe in *saa* mutants, some maturation-associated phenotypes (e.g. ROS production and bacterial killing) were elevated in *saa*^−/−^ neutrophils. Thus, Saa may differentially affect neutrophils at different stages of development and maturation.

SAA is induced in inflamed tissues, and whole animal *Saa1*^−/−^*Saa2*^−/−^ double knockout mice (which still possess *Saa3*) are sensitive to DSS-induced colitis [57], suggesting SAA protects against intestinal inflammation. Similarly, a recent report using *Saa3* knockout mice demonstrated that SAA3 suppresses bone marrow derived dendritic cell response to LPS [81], mirroring the anti-inflammatory effects of Saa on zebrafish neutrophils demonstrated here. Future work is needed to determine if Saa’s pleiotropic effects on diverse cell types are mediated by different receptors, oligomeric state, or binding other molecules [82].

Collectively, our findings highlight the importance of intestinal Saa effecting systemic neutrophil development and function, suppressing their inflammatory tone and increasing mobilization to wounds. More broadly, our findings suggest that the ontogenetic and microbial control of priming factors is important for vertebrate immunological development.

## Materials and methods

### Animal husbandry

Zebrafish studies were approved by the Institutional Animal Care and Use Committees of Duke University Medical Center (protocol number A115-16-05). All zebrafish lines were maintained on a mixed Tübingen (Tü) / TL background on a 14:10 hour light:dark cycle in a recirculating aquaculture system. From 5 dpf to 14 dpf, larvae were fed Zeigler AP100 <50-micron larval diet (Pentair, LD50-AQ) twice daily and Skretting Gemma Micro 75 (Bio-Oregon, B5676) powder once daily. Beginning at 14 dpf, larvae were fed *Artemia* (Brine Shrimp Direct, BSEACASE) twice daily, supplemented with a daily feed of Skretting Gemma Micro 75. From 28 dpf, the Gemma Micro 75 diet was replaced with Gemma Micro 300 (Bio-Oregon, B2809). At the onset of breeding age or sexual maturity, adult fish were given a 50/50 mix of Skretting Gemma Micro 500 (Bio-Oregon, B1473) and Skretting Gemma Wean 0.5 (Bio-Oregon, B2818) and one feeding of *Artemia* daily.

Larvae were also maintained on a 14:10 hour light:dark cycle in a 28.5°C incubator, and are of indeterminate sex. Gnotobiotic zebrafish were generated following natural mating and reared as method as described previously [83] with the following exception: GZM with antibiotics (AB-GZM) was supplemented with 50 μg/ml gentamycin (Sigma, G1264). Conventionally raised zebrafish were maintained at a density of ≤ 1 larva / mL, and at 3 dpf groups of 30 larvae were transferred to 10 cm petri dishes containing 20 mL gnotobiotic zebrafish media (GZM), inoculated with 3 mL filtered system water (5μm filter, SLSV025LS, Millipore) and fed autoclaved ZM-000 zebrafish diet (1% w/v stock concentration (in RO H_2_O), 0.0025% w/v final concentration, Zebrafish Management Ltd.) [10]. *TgBAC(cldn15la:EGFP)*^*pd1034Tg*^, *Tg(lyz:DsRed)*^*nz50*^ and *Tg(lyz:GFP)*^*nz117*^ have been previously characterized [40, 51].

### Zebrafish mutagenesis

Targeted deletion of the *saa* gene was performed using CRISPR/Cas9 gene editing targeting the second exon of *saa* as described [7]. Briefly, the guide RNA sequence was designed using the “CRISPR Design Tool” (http://crispr.mit.edu/). Guide RNA oligos (S1 Table, primers P1 and P2) were ligated into pT7-gRNA plasmid (Addgene, 46759), which, following BamHI (New England Biolabs, R0136L) digest, was *in vitro* transcribed using MEGAshortscript T7 kit (ThermoFisher, AM1354) [84]. Cas9 mRNA was generated from XbaI (New England Biolabs, R0145S) digested pT3TS-nls-zCas9-nls plasmid (Addgene, 46757), and *in vitro* transcribed using mMESSAGE mMACHINE T3 kit (ThermoFisher, AM1348) [84]. A cocktail consisting of 150 ng/μL of nls-zCas9-nls and 120 ng/μL of gRNA, 0.05% phenol red, 120 mM KCl, and 20 mM HEPES (pH 7.0) was prepared, and approximately 1-2 nL was injected directly into the cell of one cell stage Tü zebrafish embryos. Mutagenesis was initially screened using Melt Doctor High Resolution Melting Assay (ThermoFisher, 4409535), and identified three independent alleles which were confirmed as deletions by Sanger sequencing of TOPO-cloned PCR products. Subsequent screening of the Δ22 (allele designation *rdu60*) was performed by PCR (primers P3 and P4, S1 Table) and products were resolved on 2% agarose TBE gels. Screening of the Δ2 and Δ5 alleles (allele designations *rdu61* and *rdu62* respectively) was performed by PCR amplification using primers P3 and P4 (S1 Table), followed by purification and Hha1 digest (New England Biolabs, R0139S). All mutant alleles (*rdu60*, *rdu61*, and *rdu62*) result in the loss of a single Hha1 restriction site present in the WT sequence.

### Zebrafish transgenesis

For generation of transgenic zebrafish expressing Saa in intestinal epithelial cells (IECs), the following strategy was employed utilizing Tol2 mediated transgenesis (https://onlinelibrary.wiley.com/doi/abs/10.1002/dvdy.21343). A 349 bp region of the zebrafish *cldn15la* promoter was PCR amplified from Tü genomic DNA using primers P5 and P6 (S1 Table), digested the FesI and AscI (New England Biolabs, R0588 and R0558), and ligated into the p5E 381 vector that had been linearized with the same enzymes to generate p5E*-0.349cldn15la*. The full-length zebrafish *saa* coding sequence was PCR amplified from Tübingen whole-larvae cDNA using primers P7 and P8 (S1 Table) and subcloned into the plasmid pENTR-AleI using In Fusion (Takara Bio, 638909) to generate pME-*saa*. Both p5E-*0.349cldn15la* and pME-*saa* were verified by PCR and Sanger sequencing. A 4-way Gateway^TM^ LR reaction was performed using LR Clonase II (ThermoFisher, 12538120) to recombine p5E-*0.349cldn15la*, with pME-*saa*, and p3E-polyA 229 into pDEST 395 (which contains a bicistronic *cmlc2:EGFP* reporter), yielding the following construct: *-0.349cldn15la:saa*:*polyA*;*cmlc2:EGFP*. This plasmid was co-injected with transposase mRNA into Tü embryos at the single cell stage as described [53]. Injected F_0_ larvae were subsequently screened for mosaic EGFP expression in the heart, raised to adulthood, and used to establish stable lines for four independent alleles of *Tg*(*-0.349cldn15la*:*saa*;*cmlc2:EGFP*) (*rdu63*, *rdu64*, *rdu67*, and *rdu68*). Experiments were conducted with both larval and adult zebrafish positive for *rdu64*, *rdu67*, and *rdu68* and non-transgenic siblings used as controls.

To confirm IEC specificity of the 349 bp *cldn15la* promoter fragment, we also recombined p5E-*cldn15la*, pME-*mCherry* 386, and p3E-polyA with pDEST 394, and injected this construct into single cell Tü embryos. Mosaic F_0_ larvae were raised to adulthood, screened for germline transmission, and used to establish stable lines for three independent alleles of *Tg(−0.349cldn15la:mCherry:polyA)*. All three alleles displayed mCherry expression restricted to the intestine (not shown), and allele *rdu65* was maintained for further analysis.

### Caudal fin amputation assay

Larval zebrafish were anaesthetized with 0.75 mM Tricaine and mounted in 3% (w/v) methylcellulose (in GZM). Caudal fins were amputated using a surgical scalpel (Surgical Specialties Sharpoint, 72-2201) and fish were revived into 10 cm dishes containing either GZM (for experiments with conventionally reared larvae) or AB-GZM without gentamycin (for experiments with gnotobiotic larvae). At 15 minutes, 3 hours, and 6 hours post wounding, animals were euthanized and fixed in 4% PFA / 1× PBS overnight at 4°C on an orbital platform. Larvae were subsequently washed with 1× PBS 3-5 times at room temperature. Imaging was performed with a Leica M205 FA equipped with a Leica DFC 365FX camera using either GFP or mCherry filters. Lyz^+^ cells were enumerated at the tail wound margin using the Fiji Cell Counter plugin. Fish that were wounded at the notochord or were moribund were excluded from analysis.

### *In vivo* imaging and cell tracking analysis

Larval zebrafish were anaesthetized and mounted in 100 μL 0.75% (w/v in GZM) low melt agarose (Fisher Scientific, BP165-25) with 0.6 mM Tricaine in 96-well clear-bottom black-walled plates (Greiner Bio-One, 655090), and overlaid with 100 μl GZM containing 0.375 mM Tricaine. For homeostatic behavioral analysis, time lapse imaging was performed for 10 or 15 minutes and frames acquired at 30s intervals on a Zeiss Axio Observer with a Photometrics Evolve EMCCD camera and a 5× objective (NA 0.16, WD 18.5 mm). For live imaging following caudal fin amputation, larvae were mounted as described above and imaged for 6 hours at 2 or 5 minute intervals on an inverted Zeiss Axio Observer Z1 microscope equipped with an Xcite 120Q light source (Lumen Dynamics), an MRm camera (Zeiss), and a 20X objective lens (NA 0.4, WD 7.9 mm). Fish that were wounded at the notochord, moribund, or damaged during mounting were excluded from analysis. Automated cell tracking was performed using the MTrack2 plugin for Fiji. For cell tracking following caudal fin amputation, a region of interest (ROI) was drawn from the posterior end of the notochord to the wound margin. For neutrophil behavior analysis in homeostasis, two distinct ROIs were analyzed. An ROI (396 pixels by 52 pixels, w × h) was drawn over the intestine ending at the cloaca, and another ROI (364 pixels by 152 pixels, w × h) was positioned dorsally to the intestine in the trunk. We subsequently filtered tracking results to include cells that were tracked for ≥ 3 consecutive frames. Velocity and meandering index were calculated as previously described [12].

### Epifluorescence stereomicroscopy

Larval zebrafish from pooled clutches were anaesthetized in 0.75 mM Tricaine and mounted in 3% methylcellulose. Between 15 and 30 fish / genotype were imaged on a Leica M205 FA microscope equipped with a Leica DFC 365FX camera using identical magnification and exposures. Neutrophil recruitment to the intestine was quantified from 8-bit images with Fiji software using the Cell Counter plug in as described previously [12].

### Fluorescence Activated Cell Sorting (FACS)

For FACS, replicate pools of 60-90 lyz^+^ larvae of the indicated genotypes were euthanized with 3 mM Tricaine and washed for 5 minutes with deyolking buffer (55 mM NaCl, 1.8 mM KCl and 1.25 mM NaHCO_3_). Larvae were transferred to gentleMACS “C” tubes (Miltenyi Biotec, 130-096-334) containing 5 mL Buffer 1 [HBSS supplemented with 5% heat-inactivated fetal bovine serum (HI-FBS, Sigma, F2442) and 10 mM HEPES (Gibco, 15630-080)]. Larvae were dissociated using a combination of enzymatic and mechanical disruption. Following addition of Liberase (Roche, 05 401 119 001, 5 μg/mL final), DNaseI (Sigma, D4513, 2 μg/mL final), Hyaluronidase (Sigma, H3506, 6 U/mL final) and Collagenase XI (Sigma, C7657, 12.5 U/mL final), samples were dissociated using pre-set program C_01 on a gentleMACS dissociator (Miltenyi Biotec, 130-093-235), then incubated at 30°C on an orbital platform at 75 RPM for 10 minutes. The disruption-incubation process was repeated 4-6 times, after which 400 μL of ice-cold 120 mM EDTA (in 1× PBS) was added to each sample. Following addition of 10 mL Buffer 2 [HBSS supplemented with 5% HI-FBS, 10 mM HEPES and 2 mM EDTA], samples were filtered through 30 μm cell strainers (Miltenyi Biotec, 130-098-458) into 50 mL conical tubes.

Filters were washed with 10 mL Buffer 2, and samples were centrifuged at 1800 rcf for 15 minutes at room temperature. The supernatant was decanted, and cell pellets were resuspended in 500 μl Buffer 2. For CellROX (Invitrogen C10491 or C10444) labeling experiments, cells were then resuspended in 500 μL Buffer 2 and transferred to individual wells of a 24-well plate. CellROX was added to a final concentration of 1 μM, and samples were incubated for 45 minutes in the dark at 28.5°C on a tilting platform. Samples were transferred to FACS tubes (Falcon, 352052), and DNaseI (5 μg/mL; Sigma, D4513) and 7-AAD (Sigma, A9400, 5 μg/mL) were added. FACS was performed with either a Beckman Coulter MoFlo XDP, Beckman Coulter Astrios, or a Becton Dickinson Diva at the Duke Cancer Institute Flow Cytometry Shared Resource. Single-color control samples were used for compensation and gating. Viable neutrophils were identified as 7-AAD^−^ lyz^+^. Data were analyzed with FloJo v10 (Treestar, CA).

### *Ex vivo* zebrafish neutrophil culture

Kidneys were dissected from adult male and female transgenic lyz^+^ zebrafish of the indicated genotypes. Fish were of standard length 25.72mm±1.18mm (mean±S.D.). Single cell suspensions were generated by enzymatic treatment of dissected kidneys with DNaseI (2 μg/mL) and Liberase (5 μg/mL final) with gentle agitation on a fixed speed orbital platform (VWR, 82007-202) for 20 minutes at room temperature. Enzymes were deactivated by the addition of EDTA as described above and cell suspensions were filtered through 30 μm filters and either stained with 1 μM CellROX for 1 hour at 28°C and/or stained with 7-AAD (5 μg/mL) or Propidium Iodide (PI, Sigma, P21493, 5 μg/mL). Viable (7-AAD-or PI-) lyz^+^ cells were collected into poly-D-lysine (Sigma, P7280-5MG, 33 μg/mL) coated 96 well black-wall clear-bottom plates (Corning, 3603) containing 100 μl RPMI1640 (Gibco, 11835030) supplemented with 10% HI-FBS at a density of 15,000 cells per well. A total of 6-9 kidneys per genotype were pooled and used to seed 4-6 wells. DsRed-expressing *Escherichia coli* MG1655 (pRZT3, [85] or GFP-expressing *Pseudomonas aeruginosa* PAO1 (pMF230, [86] were cultured aerobically overnight shaking at 37°C in LB supplemented with either tetracycline (10 μg/mL) or carbenicillin (100 μg/mL), respectively. Bacteria (100 μL) were subsequently sub-cultured into 5 mL selective LB media and grown at 37°C with shaking under aerobic conditions to an OD_600_ of 0.7 - 1, diluted in sterile 1× PBS (Gibco, 14190) to a concentration of 10^4^ bacterial / μL (*E. coli*) or 10^3^ bacteria / μL (*P. aeruginosa*), and added isolated neutrophils at an MOI of or 0.2 respectively. Neutrophils were co-cultured with bacteria for 2 to 4 hours at 28°C with gentle agitation in the dark. Serial dilutions of co-culture supernatants were prepared in sterile 1× PBS and plated on selective media. Plates were incubated aerobically at 28°C for 24 hours and CFUs were enumerated.

For imaging studies, neutrophils were collected as described above with the following modification: cells were collected into poly-D-lysine coated thin-bottom 96 well plates (Greiner, 655090). Neutrophils were imaged on a Zeiss 710 inverted confocal microscope with 10× (NA 0.45) or 63× oil objectives (NA 1.40) with or without addition of bacteria.

To measure neutrophil viability, 96 well black wall clear bottom plates containing 15,000 cells / well were incubated with or without *E. coli* MG1655 (MOI 2, grown as described above) for 3.5 hours. Propidium Iodide (PI) was added to a final concentration of 6 μg/mL to each well, and plates were incubated for an additional 30 minutes at 28.5°C before reading fluorescence at 535/620 (Ex/Em, nm) with a BioTek Synergy2 plate reader. As a positive control, lyz^+^ cells were incubated with Triton X-100 (20% v/v) for 3 hours, then incubated at 65°C for 30 minutes prior to PI staining. Data are shown as % of maximum PI signal.

### Cytospins and scoring of neutrophil maturation

For morphological assessment, 15,000-30,000 viable lyz^+^ cells isolated by FACS from adult dissected kidneys as described above were sorted into 500 μl Buffer 2. Cytospins were performed immediately following collection with a Cytospin 3 (Shandon) by centrifuging cell suspensions for 3 minutes at 800 rcf. Slides were dried overnight at room temperature, fixed with absolute methanol, then stained with Wright Giemsa (Sigma, WG16) according to the manufacturer’s instructions. Slides were imaged with a Leica DMRA2 compound microscope with a PL APO 40× air objective (NA 0.85, WD 0.11mm) and Q Imaging Micropublisher Digital color camera. Twenty individual ROIs were imaged per genotype, and cells were classified based on distinct nuclear morphology by a blinded investigator as described [42].

### Gene expression analysis

Pools of 20-30 whole larvae, dissected digestive tracts, or carcasses were collected into 1 mL of TRIzol (ThermoFisher, 15596026) and stored at −80°C. Tissues were homogenized by passing samples 10-15 times through a 27-gauge needle. RNA was isolated following the manufacturer’s protocol with the following modification: a second wash with 70% ethanol (prepared in DEPC-treated H_2_O) was performed. For gene expression analysis of sorted cells, viable lyz^+^ cells were collected into 750 μL TRIzol LS (ThermoFisher 10296010). RNA was isolated using a NORGEN RNA Cleanup and Concentrator Micro-Elute Kit (Norgen Biotek, 61000), and samples eluted in 10 μL from which 8 μL of RNA was treated with DNaseI (New England Biolabs, M0303L) prior to cDNA synthesis. cDNA was synthesized using the iScript kit (Bio-Rad, 1708891). Quantitative PCR was performed in duplicate 25μl reactions using 2X SYBR Green SuperMix (PerfeCTa, Hi Rox, Quanta Biosciences, 95055) run on an ABI Step One Plus qPCR instrument using gene specific primers (S1 Table). Data were analyzed with the *ΔΔCt* method.

### *In vivo* bacterial infections

#### Pseudomonas aeruginosa

(PAO1) carrying a constitutively expressed GFP plasmid (pMF230) [86] was grown in LB media supplemented with 100 μg/mL carbenicillin overnight shaking at 37°C. Overnight culture was concentrated to an OD600 of 5 (approximately 2×10^9^ bacteria / mL) then frozen in aliquots. At 1 dpf, larvae were treated with 45 μg/mL 1-phenyl-2-thiourea (PTU) to inhibit melanization. Larvae were infected at 2 dpf by a genotype-blinded investigator. Bacteria were injected into the caudal vein with borosilicate needles along with a phenol red tracer (3% w/v) as described previously [87], and any larvae that were damaged by injections were excluded from analysis. Approximately 150-300 CFU of *P. aeruginosa* PAO1 GFP was injected per larvae. To enumerate CFUs in the inoculum, the injection dose was plated before and after infections on LB agar plates supplemented with 200ug/mL carbenicllin. Immediately following infection larvae were screened for even dosing by fluorescence microscopy, and significantly under-or over-infected larvae were excluded from further analysis by an investigator blinded to genotype. To quantify bacterial burden, individual larvae were homogenized in 500 μL sterile 1× PBS using a Tissue-Tearor (BioSpec Products, 985370) at 24 hour intervals, and serial dilutions were plated on selective media (LB agar supplemented with 100 μg/mL carbenicillin). *P. aeruginosa* CFUs were enumerated following aerobic incubation for 24 hours at 28°C.

### Zebrafish immunohistochemistry

For immunostaining, 6 dpf larval zebrafish were fixed in 4% PFA/1× PBS overnight at 4°C on an orbital platform, then washed 3-5× with 1× PBS the following day. Larvae were mounted in 4% low melt agarose in cryo-molds molds (Tissue-Tek) and 200 μm transverse sections cut using a Leica VT1000S vibratome. Sections were transferred to 24-well plates, washed 3× in 1× PBS at room temperature, and permeabilized with 1× with PBS containing 0.1% (v/v) Triton ×-100 (PBS-T) for 30 minutes at RT. Sections were blocked in 5% (v/v) donkey serum in PBS-T for 1 hour at room temperature, then incubated with primary antibodies diluted in blocking buffer overnight at 4°C with agitation (Rabbit anti 4E8, Abcam, ab73643, 1:200; Rabbit anti DsRed, Clontech, 632496, 1:200; Chicken anti GFP, AVES, GFP-10×0, 1:200). Sections were washed with PBS, then incubated with species-specific secondary antibodies diluted in PBS-T for 4 hours at room temperature on a tilting platform [(ThermoFisher, A10042, A32728, A11039, 1:200) and Hoechst 33258 (ThermoFisher, H3569, 1:1000)]. Sections were then washed 3× with 1× PBS then mounted and coverslipped on slides using mounting media containing DAPI (Vector Laboratories, Inc, H-1200). Slides were imaged with a Zeiss LSM 780 upright confocal microscope equipped with a GaAsP array detector using a 63× oil objective (NA 1.4, WD 0.19 mm).

### 16S rRNA gene sequencing

Adult heterozygous *saa*^*rdu60*/+^ zebrafish were bred naturally and the resulting embryos were collected into system water and pooled at 0 dpf. Fertilized embryos were sorted at equal densities into autoclaved 3L tanks filled with system water at 1 dpf (50 embryos per 3 liter tank). Larvae were maintained under static conditions until 6 dpf, at which time water flow and feeding were begun. Dissected digestive tracts and environmental water samples were collected at two time points: 6 dpf and 70 dpf. At 6 dpf zebrafish were sampled prior to first feeding, and fish sampled at 70 dpf were fed using the standard facility diet (as described above in animal husbandry) beginning at 6 dpf. For water samples, 50 mL of water was collected from each tank and filtered using 0.2μm MicroFunnel Filter Units (Pall Corporation, 4803). Filters were removed from filter units with sterile forceps, transferred to Eppendorf tubes and snap frozen in a dry ice/ethanol bath. For intestinal samples, digestive tracts were dissected and flash frozen and either carcasses or fin were reserved for genotyping. Fish samples were genotyped to identify homozygous *saa*^+/+^ and *saa*^*rdu60/rdu60*^ zebrafish prior to submission for genomic DNA extraction.

The Duke Microbiome Shared Resource (MSR) extracted bacterial DNA from gut and water samples using a MagAttract PowerSoil^®^ DNA EP Kit (Qiagen, 27100-4-EP) that allows for the isolation of samples in a 96 well plate format using a Retsch MM400 plate shaker. DNA was extracted from ≥12 fish per genotype per time point, and from 5 to 8 different tanks per timepoint to control for tank effects. Sample DNA concentration was assessed using a Qubit dsDNA HS assay kit (ThermoFisher, Q32854) and a PerkinElmer Victor plate reader. Bacterial community composition in isolated DNA samples was characterized by amplification of the V4 variable region of the 16S rRNA gene by polymerase chain reaction using the forward primer 515 and reverse primer 806 following the Earth Microbiome Project protocol (http://www.earthmicrobiome.org/). These primers (515F and 806R) carry unique barcodes that allow for multiplexed sequencing. Equimolar 16S rRNA PCR products from all samples were quantified and pooled prior to sequencing. Sequencing was performed by the Duke Sequencing and Genomic Technologies shared resource on an Illumina MiSeq instrument configured for 150 base-pair paired-end sequencing runs.

Subsequent data analysis was conducted in QIIME2 (https://qiime2.org), the successor of QIIME [88]. Paired reads were demultiplexed with qiime demux emp-paired, and denoised with qiime dada2 denoise-paired [89]. Taxonomy was assigned with qiime feature-classifier classify-sklearn [90], using a naive Bayes classifier, trained against the 99% clustered 16S reference sequence set of SILVA, v. 1.19 [91]. A basic statistical diversity analysis was performed, using qiime diversity core-metrics-phylogenetic, including alpha-and beta-diversity, as well as relative taxa abundances in sample groups. The determined relative taxa abundances were further analyzed with LEfSe (Linear discriminant analysis effect size) [92], to identify differential biomarkers in sample groups.

### Statistical methods

All experiments were repeated at least two times and statistical analyses were performed with GraphPad Prism v.7. Data are presented as mean ± SEM. For comparisons between 2 groups a two tailed student’s *t*-test or Mann-Whitney test was applied. For comparisons between 3 or more groups, a one-way ANOVA with Tukey’s multiple comparisons test was used. For experiments with 2 independent variables, a two-way ANOVA was performed. Significance was set *as* p < 0.05, and denoted as: * *p* < 0.05, ** *p* < 0.01, *** *p* < 0.001, **** *p* < 0.0001. Sample sizes are indicated in the figure legends.

### Data availability

All data are available from authors upon request. 16S rRNA sequencing data have been deposited in the NCBI SRA (accession number pending).

## Acknowledgements

The authors are grateful to Nancy Martin, and Bin Li, PhD and Mike Cook, PhD (Duke Cancer Institute Flow Cytometry Resource) for their assistance with FACS, and Lisa Cameron, PhD, Ben Carlson, PhD and Yasheng Gao, PhD (Duke Light Microscopy Core Facility) for their assistance with instrumentation. We are grateful to Holly Dressman, PhD and Zhengzheng Wei of the Duke Microbiome Shared Resource for DNA isolation and 16S rRNA gene sequencing. We are also grateful to Colin Lickwar, PhD for assistance and advice with bioinformatics. This work was supported by the NIH (P01-DK094779) awarded to J.F.R. and AI130236 (D.M.T.). C.C.M. was supported by the US National Science Foundation GRFP fellowship (DGE-1644868).

## Author Contributions

Conceptualization, C.C.M. and J.F.R. Methodology, C.C.M. and S.T.E. Investigation, C.C.M., S.T.E., and M.A.M. Formal Analysis, C.C.M., S.T.E., O.M. Writing Original Draft, C.C.M. and J.F.R. Writing-Review and Editing, C.C.M., J.F.R., S.T.E., M.A.M., D.M.T. Data curation, O.M. Funding acquisition, C.C.M, D.M.T., and J.F.R. Supervision, D.M.T. and J.F.R.

## Supporting information

**S1 Fig.**
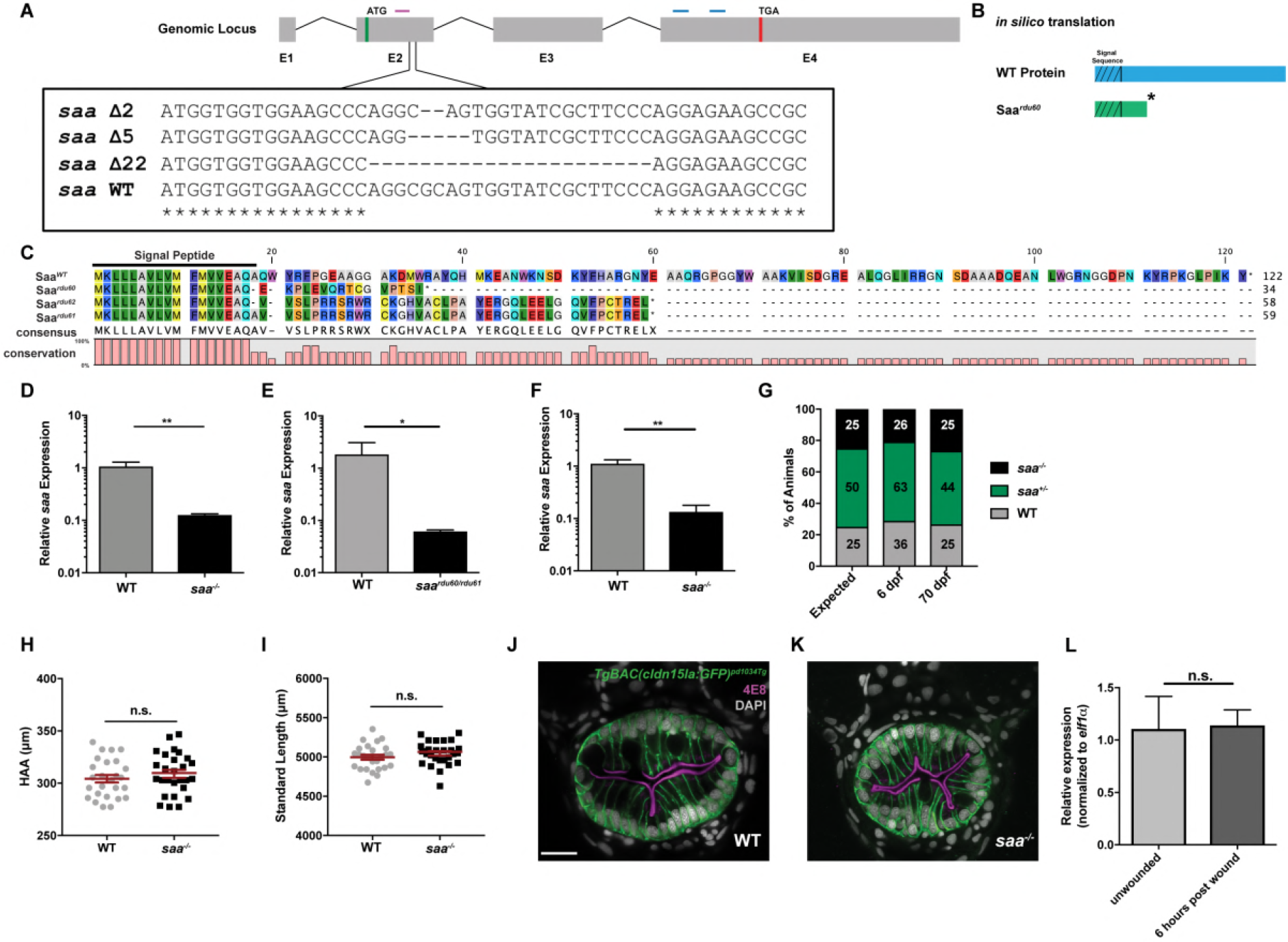
Generation of *saa* mutant zebrafish. **(A)** Sequence of CRISPR/Cas9 induced deletions in the coding region of zebrafish *saa* exon 2. **(B-C)** *In silico* translation of mutant allele *rdu60* shows a frame shift mutation in exon 2 with a predicted early stop codon, indicated by the asterisk. **(D)** qRT-PCR of 6 dpf whole larvae shows *saa* homozygous mutant zebrafish (*saa*^*rdu60/rdu60*^ or *saa*^−/−^) express negligible levels of *saa* mRNA. **(E)** qRT-PCR of 6 dpf whole larvae from trans-heterozygous *saardu60/rdu61* in-crosses demonstrate reduced *saa* mRNA levels relative to WT controls. **(F)** qRT-PCR of dissected digestive tracts from 6 dpf WT and *saa*^−/−^ larvae show significantly reduced *saa* mRNA levels (qRT-PCR shown in panels B-D included 4-8 replicates / genotype, n ≥ 20 larvae / replicate). **(G)** Representative genotype distributions show there is no deviation from expected Mendelian outcomes indicating no difference in viability of *saa*^−/−^ animals (*p =* 0.8257) (from two independent experiments, actual number of animals from each genotype overlaid on bars). **(H-I)** Morphometric analysis shows loss of *saa* does not impact growth [standard length (SL), height at anterior of anal fin (HAA)] in 6 dpf larvae (n ≥ 26 larvae). **(J-K)** Representative immunofluorescence images of from 200 μm thick sections of *TgBAC(cldn15la:EGFP)*^*pd1034Tg*^ WT and *saa*^−/−^ larvae stained with the brush-border antibody 4E8 demonstrate intestinal architecture is qualitatively normal in mutant animals (scale bar = 50 μm). **(L)** qRT-PCR of 6 dpf whole larvae following a tail amputation shows no significant induction of *saa.* In panels D, E, F, H, I, and L a *t*-test was used. In panel E, a Mann-Whitney test was applied. A Chi-squared test was used to test for deviation from Mendelian distribution in G. Data are presented as mean ± SEM. * *p* < 0.05, ** *p* < 0.01, *** *p* < 0.001, **** *p* < 0.0001

**S2 Fig.**
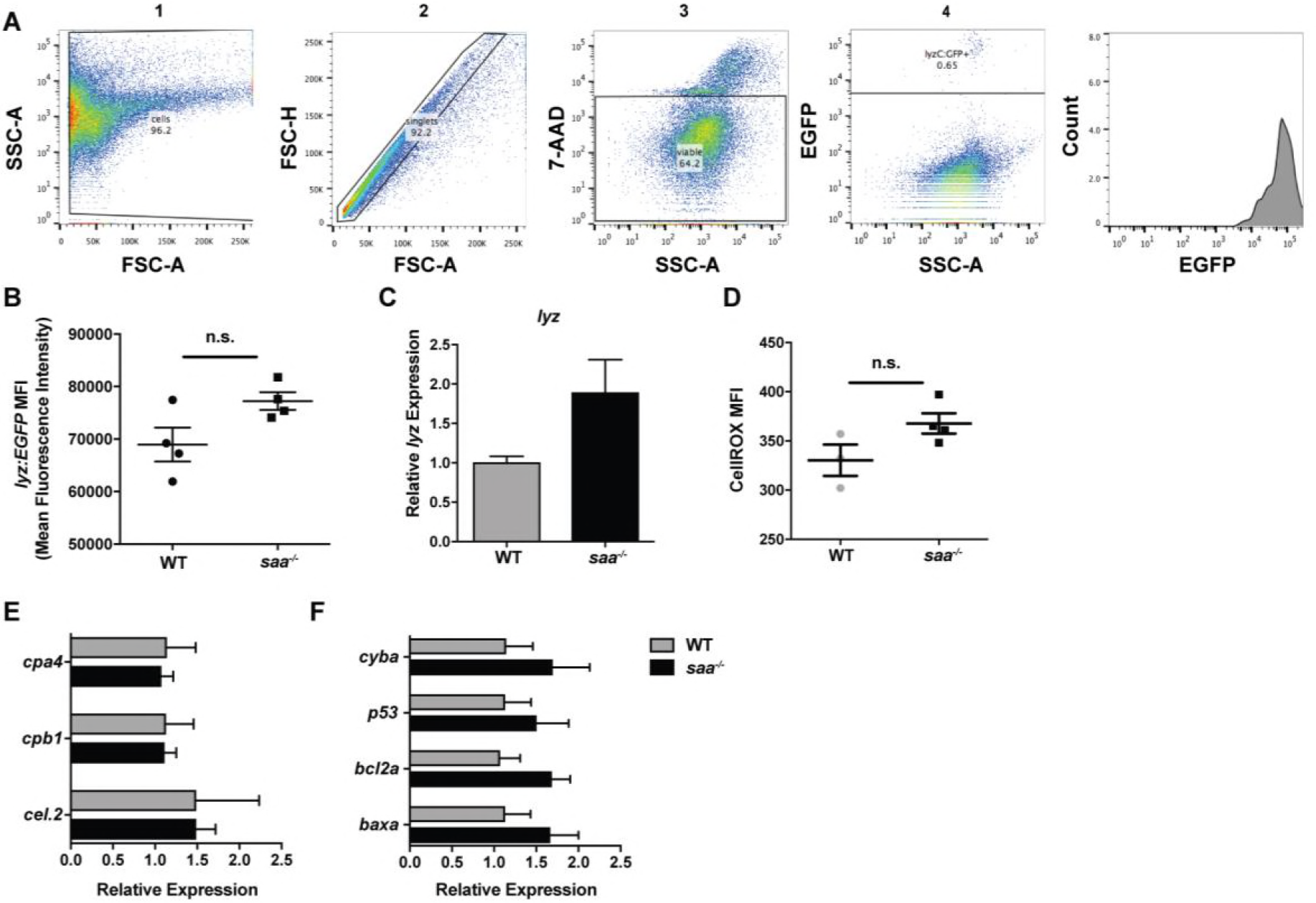
Isolation and characterization of neutrophils from WT and *saa* mutant zebrafish larvae. **(A)** Gating strategy for isolation of lyz:EGFP^+^ neutrophils from 6 dpf zebrafish larvae. **(B)** The mean fluorescence intensity (MFI) of the lyz:EGFP^+^ neutrophil population is not significantly different between WT and *saa* mutant larvae **(C)** qRT-PCR shows no significant difference in *lysozyme C* (*lyz*) transcript levels in sorted lyz^+^ neutrophils from WT and *saa* mutant larvae (*p* = 0.0789). **(D)** Quantification of intracellular ROS levels as indicated by CellROX staining measured by flow cytometry in lyz:EGFP^+^ neutrophils from WT and *saa* mutant larvae shows no significant difference. **(E-F)** qRT-PCR analysis of sorted neutrophils reveals no differential expression of genes associated with pro-myelocyte progenitors (*cpa4*, *cpb1*, *cel.2*) or apoptotic markers (*cyba, p53, bcl2a, baxa*) between WT or *saa* mutants (For panels B - F: n ≥ 4 replicates / genotype, n = 60 90 larvae / genotype). In panels B-F data was analyzed by *t*-test. Data are presented as mean ± SEM. * *p* < 0.05, ** *p* < 0.01, *** *p* < 0.001, **** *p* < 0.0001

**S3 Fig.**
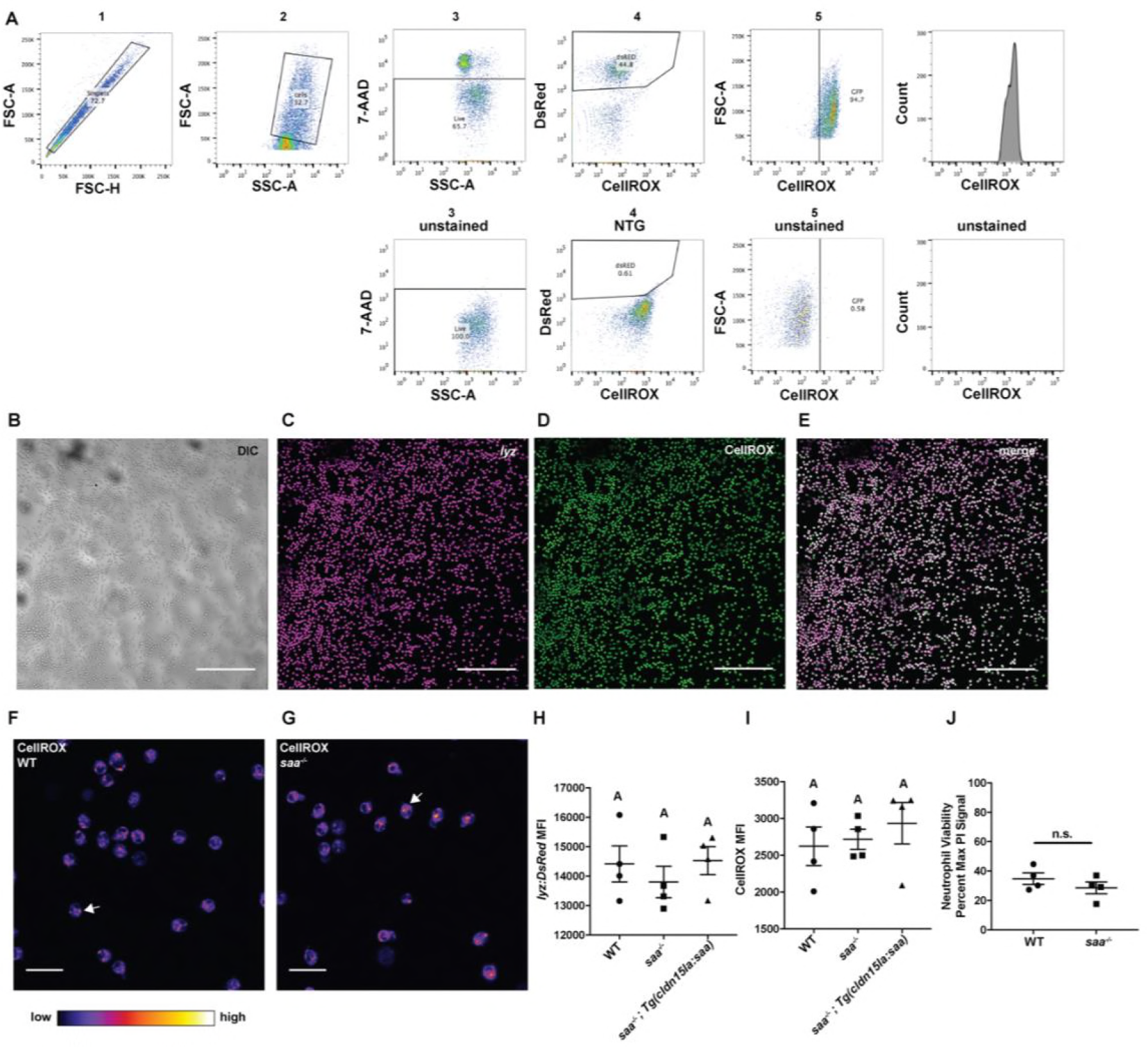
Isolation and culture of zebrafish neutrophils *ex vivo*. **(A)** Gating strategy for isolation of CellROX labeled lyz:DsRed^+^ neutrophils from adult zebrafish kidneys. **(B-E)** Low magnification (10x) confocal images of lyz:EGFP^+^ neutrophils labeled with CellROX *ex vivo* (scale bar = 200 μm). **(F-G)** Imaging of CellROX labeled lyz:EGFP^+^ neutrophils isolated from WT and *saa* mutant zebrafish reveals cytoplasmic punctae (indicated by white arrows) (Scale bar = 20 μm). Neutrophils were imaged with a Zeiss 710 inverted confocal microscope with 10× (NA 0.45) or 63× oil objectives (NA 1.40). **(H)** The mean fluorescence intensity (MFI) of the lyz:DsRed^+^ population is not significantly different between WT, *saa*^−/−^, or *saa*^−/−^;*Tg(cldn15la:saa)* neutrophils (4 replicates / genotype). **(I)** Quantification of intracellular ROS levels as indicated by CellROX staining measured by flow cytometry of lyz:DsRed^+^ neutrophils shows no significant difference in baseline CellROX levels between genotypes (4 replicates / genotype). **(J)** Measurement of lyz:EGFP^+^ neutrophil viability as assessed by Propidium Iodide (PI) staining shows no significant differences between genotypes after 4 hours of co-culture with *E. coli* (relative to maximum PI signal from lysed neutrophils) (4 replicates / genotype). In panels H-I, data were analyzed by one-way ANOVA with Tukey’s multiple comparisons test. In panel J, data were analyzed by *t*-test. Data are presented as mean ± SEM. * *p* < 0.05, ** *p* < 0.01, *** *p* < 0.001, **** *p* < 0.0001

**S4 Fig.**
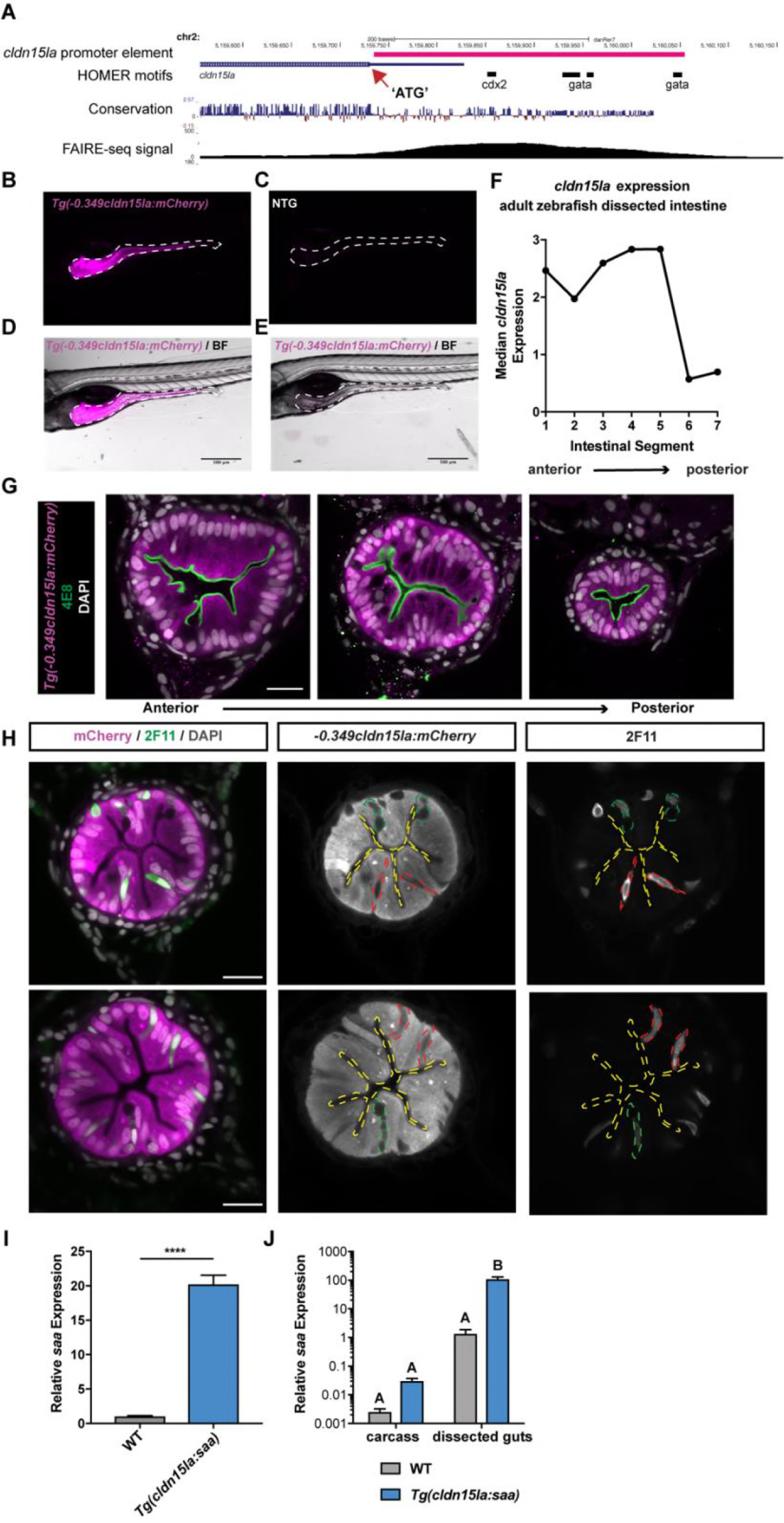
Characterization of the zebrafish *cldn15la* promoter used to drive intestine-specific *saa* expression. **(A)** UCSC genome browser view of the zebrafish *cldn15la* gene locus with the translational start indicated by the red arrow. Pink bar represents the cloned 349 bp promoter region upstream of the *cldn15la* gene used to drive intestine-specific transgene expression. Tracks for vertebrate conservation, FAIRE-seq and motifs for transcription factors important of IEC gene expression programs (identified by HOMER) are shown below the locus [53]. **(B-E)** Representative stereoscope images of IEC specific mCherry expression in 6 dpf *Tg(−0.349cldn15la:mCherry)*^*rdu65*^ larvae compared to non-transgenic (NTG) controls (scale bar = 500 μm). **(F)** Expression pattern of endogenous *cldn15la* along the length of the intestine in adult zebrafish, as measured by microarray in Wang et al., 2010 [52]. **(G)** Representative confocal micrographs of immunostained transverse sections of *Tg(−0.349cldn15la:mCherry)* larvae along the anterior-posterior axis labeled with the absorptive cell brush border-specific antibody 4E8 illustrates transgene expression in absorptive enterocytes (scale bar = 20 μm). **(H)** Representative immunofluorescence images of transverse sections from *Tg(−0.349cldn15la:mCherry)* larvae stained with secretory cell-specific antibody 2F11 demonstrates that the transgene is weakly expressed in secretory cells, including enteroendocrine cells (outlined by red dashed lines) and goblet cells (outlined by green dashed lines) (scale bar = 20 μm). **(I)** qRT-PCR of 6 dpf *Tg(cldn15la:saa)* and WT sibling whole larvae shows elevated *saa* mRNA levels in transgenic larvae (5 replicates / genotype, n = 25 larvae / replicate). Significance determined by unpaired *t*-test. **(J)** qRT-PCR analysis of dissected digestive tracts and carcasses from *Tg(cldn15la:saa)*^*rdu63*^ and WT sibling 6 dpf larvae shows significant increase in *saa* expression in transgenic larvae is restricted to the gut (5 replicates / genotype, 15 larvae / replicate). Data analyzed by one-way ANOVA with Tukey’s multiple comparison’s test. Data are presented as mean ± SEM. * *p* < 0.05, ** *p* < 0.01, *** *p* < 0.001, **** *p* < 0.0001

**S5 Fig.**
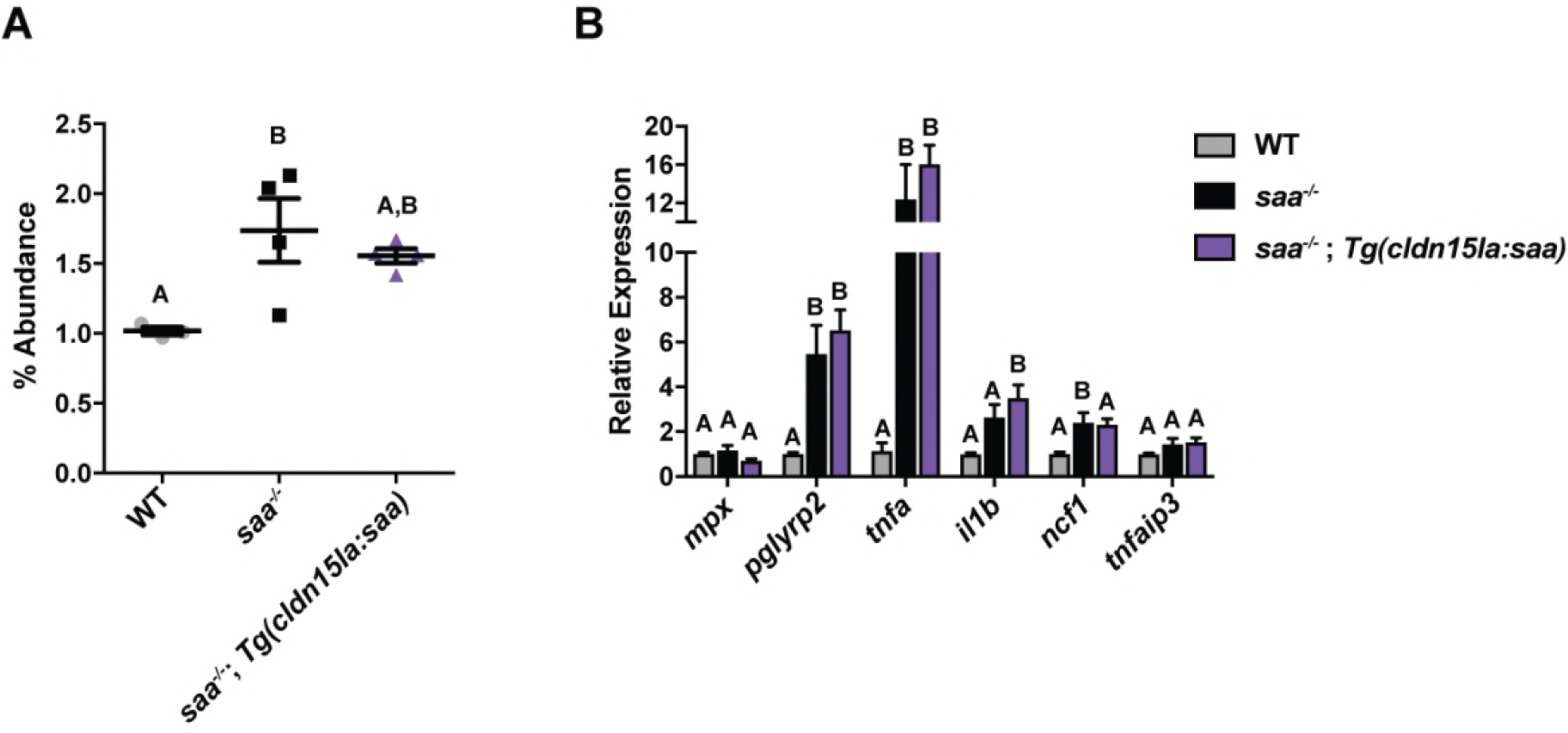
Intestinally-derived Saa is insufficient to restore elevated abundance and transcriptional activation of neutrophils in mutant zebrafish larvae. **(A)** FACS reveals increased abundance of lyz:DsRed^+^ neutrophils in 6 dpf *saa*^−/−^ larvae as well as *saa*^−/−^;*Tg(cldn15la:saa)* larvae compared to WT controls. **(B)** qRT-PCR of pro-inflammatory mRNAs from lyz:DsRed^+^ neutrophils isolated from both *saa*^−/−^ and *saa*^−/−^;*Tg(cldn15la:saa)* larvae as compared to WT (4 replicates / genotype, n ≥ 60 larvae / replicate). Data in panels A and B were analyzed by one-way ANOVA with Tukey’s multiple comparisons test. Data are presented as mean ± SEM. * *p* < 0.05, ** *p* < 0.01, *** *p* < 0.001, **** *p* < 0.0001

**S1 Table.**
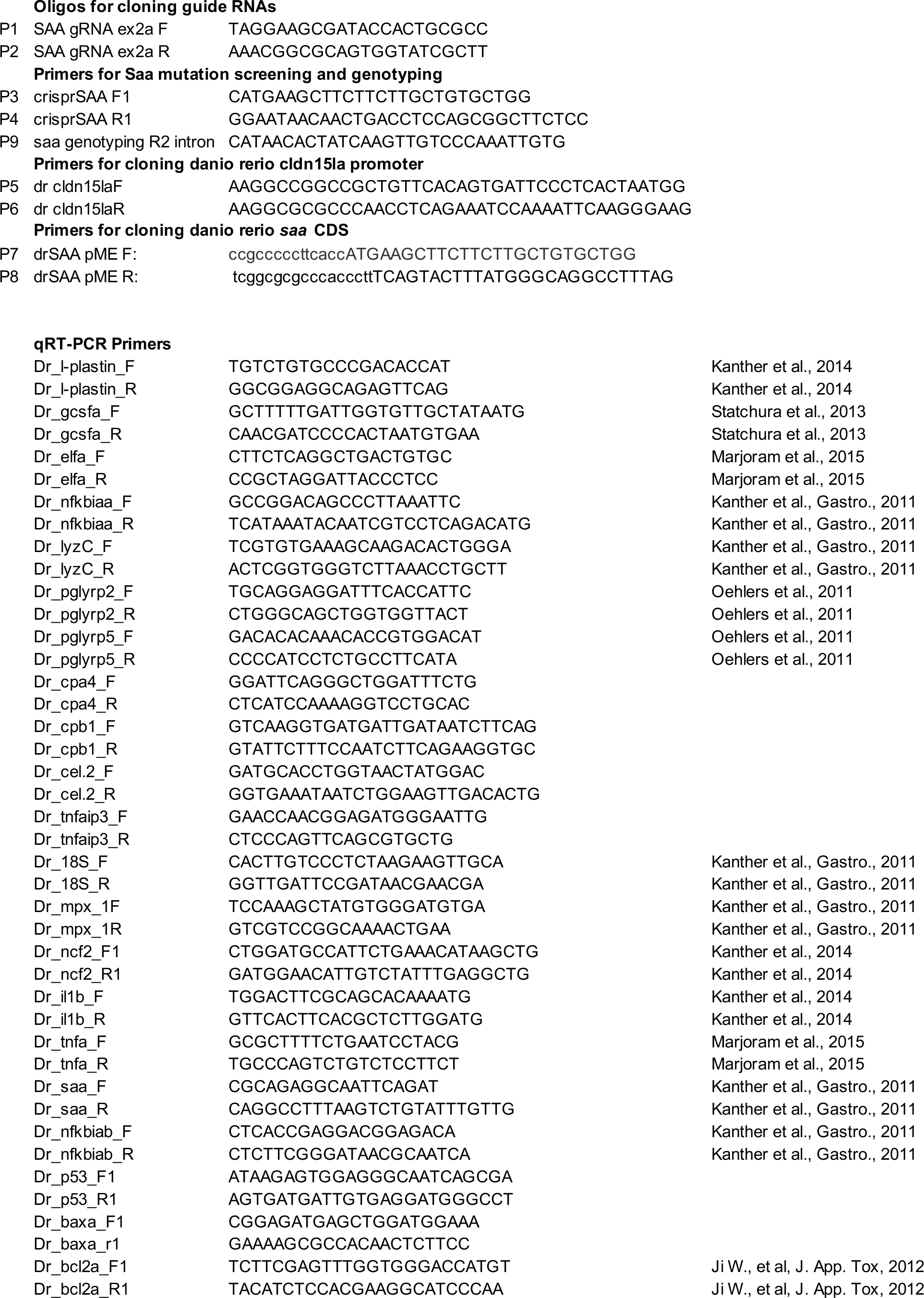
Primers used in this study (qRT-PCR, cloning and genotyping)

**S2 Table.** Statistical analysis of alpha-diversity comparisons from 16S rRNA gene sequencing (Kruskal-Wallis test) Bacterial communities in WT and *saa*^−/−^ mutant (MUT) guts were not significantly different by any of these metrics, whereas WT/MUT guts were significantly different from their housing water (H2O) as expected.

|  |  | **Chao1** |  |  |
| --- | --- | --- | --- | --- |
| **Group 1** | **Group 2** | **H** | **p-value** | **q-value** |
| 6d.H2O (n=8) | 6d.MUT (n=15) | 10.84821958 | 0.000988908 | 0.002439429 |
| 6d.H2O (n=8) | 6d.WT (n=14) | 7.835169492 | 0.005123935 | 0.006987184 |
| 6d.MUT (n=15) | 6d.WT (n=14) | 0.09340235 | 0.759895529 | 0.759895529 |
| 70d.H2O (n=5) | 70d.MUT (n=12) | 10 | 0.001565402 | 0.002609004 |
| 70d.H2O (n=5) | 70d.WT (n=15) | 10.73042169 | 0.001053884 | 0.002439429 |
| 70d.MUT (n=12) | 70d.WT (n=15) | 0.729835014 | 0.392936672 | 0.421003577 |

|  |  | **Evenness** |  |  |
| --- | --- | --- | --- | --- |
| **Group 1** | **Group 2** | **H** | **p-value** | **q-value** |
| 6d.H2O (n=8) | 6d.MUT (n=15) | 10.41666667 | 0.001248831 | 0.006020723 |
| 6d.H2O (n=8) | 6d.WT (n=14) | 7.453416149 | 0.006331617 | 0.010552694 |
| 6d.MUT (n=15) | 6d.WT (n=14) | 0.007619048 | 0.930443263 | 0.930443263 |
| 70d.H2O (n=5) | 70d.MUT (n=12) | 5.377777778 | 0.02039484 | 0.03059226 |
| 70d.H2O (n=5) | 70d.WT (n=15) | 4.954285714 | 0.026026074 | 0.035490101 |
| 70d.MUT (n=12) | 70d.WT (n=15) | 2.002380952 | 0.15705234 | 0.196315425 |

|  |  | **Faith Phylogenetic Diversity** |  |  |
| --- | --- | --- | --- | --- |
| **Group 1** | **Group 2** | **H** | **p-value** | **q-value** |
| 6d.H2O (n=8) | 6d.MUT (n=15) | 15 | 0.000107511 | 0.000403167 |
| 6d.H2O (n=8) | 6d.WT (n=14) | 13.58385093 | 0.00022814 | 0.000570349 |
| 6d.MUT (n=15) | 6d.WT (n=14) | 0.121904762 | 0.726977734 | 0.726977734 |
| 70d.H2O (n=5) | 70d.MUT (n=12) | 10 | 0.001565402 | 0.002348103 |
| 70d.H2O (n=5) | 70d.WT (n=15) | 10.71428571 | 0.001063115 | 0.001989576 |
| 70d.MUT (n=12) | 70d.WT (n=15) | 0.952380952 | 0.329113986 | 0.352622128 |

|  |  | **Observed OTUs** |  |  |
| --- | --- | --- | --- | --- |
| **Group 1** | **Group 2** | **H** | **p-value** | **q-value** |
| 6d.H2O (n=8) | 6d.MUT (n=15) | 10.84821958 | 0.000988908 | 0.002439429 |
| 6d.H2O (n=8) | 6d.WT (n=14) | 7.835169492 | 0.005123935 | 0.006987184 |
| 6d.MUT (n=15) | 6d.WT (n=14) | 0.09340235 | 0.759895529 | 0.759895529 |
| 70d.H2O (n=5) | 70d.MUT (n=12) | 10 | 0.001565402 | 0.002609004 |
| 70d.H2O (n=5) | 70d.WT (n=15) | 10.73042169 | 0.001053884 | 0.002439429 |
| 70d.MUT (n=12) | 70d.WT (n=15) | 0.729835014 | 0.392936672 | 0.421003577 |

|  |  | **Shannon** |  |  |
| --- | --- | --- | --- | --- |
| **Group 1** | **Group 2** | **H** | **p-value** | **q-value** |
| 6d.H2O (n=8) | 6d.MUT (n=15) | 5.4 | 0.020136752 | 0.037756409 |
| 6d.H2O (n=8) | 6d.WT (n=14) | 4.770186335 | 0.028956691 | 0.048261152 |
| 6d.MUT (n=15) | 6d.WT (n=14) | 0.007619048 | 0.930443263 | 0.983453831 |
| 70d.H2O (n=5) | 70d.MUT (n=12) | 8.711111111 | 0.003162764 | 0.012805467 |
| 70d.H2O (n=5) | 70d.WT (n=15) | 6.63047619 | 0.010024846 | 0.025062115 |
| 70d.MUT (n=12) | 70d.WT (n=15) | 1.60952381 | 0.204558753 | 0.278943754 |

**Simpson**
| **Group 1** | **Group 2** | **H** | **p-value** | **q-value** |
| --- | --- | --- | --- | --- |
| 6d.H2O (n=8) | 6d.MUT (n=15) | 5.4 | 0.020136752 | 0.050341879 |
| 6d.H2O (n=8) | 6d.WT (n=14) | 6.377329193 | 0.011558724 | 0.034676173 |
| 6d.MUT (n=15) | 6d.WT (n=14) | 0.007619048 | 0.930443263 | 0.930443263 |
| 70d.H2O (n=5) | 70d.MUT (n=12) | 7.511111111 | 0.006131953 | 0.022994825 |
| 70d.H2O (n=5) | 70d.WT (n=15) | 7.56 | 0.005967799 | 0.022994825 |
| 70d.MUT (n=12) | 70d.WT (n=15) | 1.866666667 | 0.171857339 | 0.214821674 |

**S3 Table.** Statistical analysis of beta-diversity comparisons from 16S rRNA gene sequencing (PERMANOVA) Bacterial communities in WT and *saa*^−/−^ mutant (MUT) guts were not significantly different by any of these metrics, whereas WT/MUT guts were significantly different from their housing water (H2O) as expected.

| **Unweighted UNIFRAC** |  |  |  |  |  |
| --- | --- | --- | --- | --- | --- |
| **Group 1** | **Group 2** | **Sample size** | **pseudo-F** | **p-value** | **q-value** |
| 6d.H2O | 6d.MUT | 23 | 9.710676626 | 0.001 | 0.001363636 |
| 6d.H2O | 6d.WT | 22 | 9.665690878 | 0.001 | 0.001363636 |
| 6d.MUT | 6d.WT | 29 | 0.916930734 | 0.553 | 0.553 |
| 70d.H2O | 70d.MUT | 17 | 10.81391064 | 0.002 | 0.002307692 |
| 70d.H2O | 70d.WT | 20 | 12.37290625 | 0.001 | 0.001363636 |
| 70d.MUT | 70d.WT | 27 | 1.275344245 | 0.119 | 0.1275 |
| **Weighted UNIFRAC** |  |  |  |  |  |
| **Group 1** | **Group 2** | **Sample size** | **pseudo-F** | **p-value** | **q-value** |
| 6d.H2O | 6d.MUT | 23 | 11.72303376 | 0.001 | 0.0015 |
| 6d.H2O | 6d.WT | 22 | 10.35076436 | 0.001 | 0.0015 |
| 6d.MUT | 6d.WT | 29 | 0.393271902 | 0.934 | 0.934 |
| 70d.H2O | 70d.MUT | 17 | 8.998130913 | 0.001 | 0.0015 |
| 70d.H2O | 70d.WT | 20 | 10.93560994 | 0.001 | 0.0015 |
| 70d.MUT | 70d.WT | 27 | 0.396353439 | 0.816 | 0.874285714 |
| **Bray-Curtis** |  |  |  |  |  |
| **Group 1** | **Group 2** | **Sample size** | **pseudo-F** | **p-value** | **q-value** |
| 6d.H2O | 6d.MUT | 23 | 8.191011555 | 0.001 | 0.001153846 |
| 6d.H2O | 6d.WT | 22 | 8.314089531 | 0.001 | 0.001153846 |
| 6d.MUT | 6d.WT | 29 | 0.792868311 | 0.819 | 0.819 |
| 70d.H2O | 70d.MUT | 17 | 6.29605003 | 0.001 | 0.001153846 |
| 70d.H2O | 70d.WT | 20 | 7.071145881 | 0.001 | 0.001153846 |
| 70d.MUT | 70d.WT | 27 | 0.449691029 | 0.801 | 0.819 |
| **Jaccard** |  |  |  |  |  |
| **Group 1** | **Group 2** | **Sample size** | **pseudo-F** | **p-value** | **q-value** |
| 6d.H2O | 6d.MUT | 23 | 5.264493744 | 0.001 | 0.00125 |
| 6d.H2O | 6d.WT | 22 | 5.400166784 | 0.001 | 0.00125 |
| 6d.MUT | 6d.WT | 29 | 0.916218743 | 0.661 | 0.661 |
| 70d.H2O | 70d.MUT | 17 | 7.533691 | 0.001 | 0.00125 |
| 70d.H2O | 70d.WT | 20 | 7.692192669 | 0.001 | 0.00125 |
| 70d.MUT | 70d.WT | 27 | 1.174809096 | 0.217 | 0.2325 |

